# A fast linkage disequilibrium-based statistical test for Genome-Wide Epistatic Selection Scans in structured populations

**DOI:** 10.1101/2020.02.14.949206

**Authors:** Léa Boyrie, Corentin Moreau, Florian Frugier, Christophe Jacquet, Maxime Bonhomme

## Abstract

The quest for genome-wide signatures of selection in populations using SNP data has proven efficient to uncover genes involved in conserved or adaptive molecular functions, but none of the statistical methods were designed to identify interacting genes as targets of selective processes. Here, we propose a straightforward statistical test aimed at detecting epistatic selection, based on a linkage disequilibrium (LD) measure accounting for population structure and heterogeneous relatedness between individuals. SNP-based (*T*_*rv*_) and window-based (*T*_*corPC*1_*_v_*) statistics fit a Student distribution, allowing to easily and quickly test the significance of correlation coefficients in the frame of Genome-Wide Epistatic Selection Scans (GWESS) using candidate genes as baits. As a proof of concept, use of SNP data from the *Medicago truncatula* symbiotic legume plant uncovered a previously unknown gene coadaptation between the *MtSUNN* (*Super Numeric Nodule*) receptor and the *MtCLE02* (*CLAVATA3-Like*) signalling peptide, and experimental evidence accordingly supported a *MtSUNN*-dependent negative role of *MtCLE02* in symbiotic root nodulation. Using human HGDP-CEPH SNP data, our new statistical test uncovered strong LD between *SLC24A5* and *EDAR* worldwide, which persists after correction for population structure and relatedness in Central South Asian populations. This result suggests adaptive genetic interaction or coselection between skin pigmentation and the ectodysplasin pathway involved in the development of ectodermal organs (hairs, teeth, sweat glands), in some human populations. Applying this approach to genome-wide SNP data will foster the identification of evolutionary coadapted gene networks.

**Author summary:** Population genomic methods have allowed to identify many genes associated with adaptive processes in populations with complex histories. However, they are not designed to identify gene coadaptation between genes through epistatic selection, in structured populations. To tackle this problem, we developed a straightforward LD-based statistical test accounting for population structure and heterogeneous relatedness between individuals, using SNP-based (*T*_*rv*_) or windows-based (*T*_*corPC*1*v*_) statistics. This allows easily and quickly testing for significance of correlation coefficients between polymorphic loci in the frame of Genome Wide Epistatic Selection Scans (GWESS). Following detection of gene coadaptation using SNP data from human and the model plant *Medicago truncatula*, we report experimental evidence of genetic interaction between two receptors involved in the regulation of root nodule symbiosis in *Medicago truncatula*. This test opens new avenues for exploring the evolution of genes as interacting units and thus paves the way to infer new networks based on evolutionary coadaptation between genes.

## Introduction

Genes or genomic regions targeted by natural selection are involved in population adaptive responses to changing environments or in the conservation of pivotal molecular functions. Patterns of DNA sequence polymorphisms in these regions may reflect signatures of directional or balancing, positive selection on adaptive mutations, or negative/background selection against deleterious mutations (1–3). Identifying genes showing such selection signatures has been a major goal of population genetics over the last decades. A large number of statistical methods have been developed, with the aim of controlling at best the false positive proportion and increasing detection power, by taking into account the neutral evolution expected for molecular polymorphisms in populations showing diverse population structures or particular demographic scenarios (2, 4, 5). Thanks to high-throughput sequencing technologies, these methods can now be used to perform Genome-Wide Selection Scans (GWSS) using massive Single Nucleotide Polymorphism (SNP) datasets (6–8). Although GWSS have identified cohorts of genes associated with past or ongoing adaptive processes, they are not designed to identify gene coadaptation, that is the adaptive interactions between genes through epistatic selection, because the genetic background on which adaptive mutations occur is not considered in statistical frameworks (9).

Few theoretical studies have addressed the issue of simulating the behavior and detecting epistatic selection in population samples using DNA polymorphisms. Simulations of two-locus epistatic models with different degrees of recombination (i.e. 0 < *c* < 0.5) in a panmictic population as led to the conclusion that, contrary to *de novo* coselected mutations or mutations disconnected by a certain amount of time before being coselected, standing variation on neutral or preselected mutations was crucial for the efficacy of epistatic selection relative to genetic drift, and therefore for its statistical detection (10, 11). Simulations of two-locus coadaptation in subdivided populations have also shown that moderate migration rates allowed the propagation of coselected mutations across subpopulations, while at the same time preserving favorable allelic combinations established within each subpopulation, thus favoring the ultimate fixation of coadapted haplotypes (12).

Adaptive epistatic interactions between alleles at two independent loci are expected to generate Linkage Disequilibrium (LD). It has been shown that a classical LD measure, such as the correlation coefficient *r* (related to *r^2^*), can be used to detect epistatic selection between two bi-allelic loci in a population because it is a directional measure which can indicate an excess of ancestral and derived allelic associations, relative to recombinant allelic associations (13). In the case where the ancestral and derived alleles at both loci are known, authors show that this measure detects epistatic selection either in a coadaptation model where two derived alleles can form a coadapted allelic combination, or in a compensatory model where the two derived alleles are individually deleterious but compensate when combined. More recently, based on simulations of structured populations of which some diverged through epistatic selection, the D’_IS_^2^ measure of LD – originally described by Ohta (14, 15) – was proposed to detect epistatic selection (16). However, this statistic showed strong limitations to distinguish between epistatic selection and single-locus selection at two independent loci. In addition, neutral forces such as population structure, genetic drift and relatedness among individuals, due to non-random mating schemes act as confounders because they naturally tend to increase genome-wide levels of LD and to generate long-distance LD (17–20), which can falsely be detected as signatures of epistatic selection (21).

A significant improvement towards capturing more specifically the LD due to physical linkage in a population with various degrees of relatedness among individuals, was the introduction of *r^2^_v_*, a *r^2^* measure. This measure includes a kinship matrix to capture relatedness among individuals in structured populations (18). Subsequently, to identify interspecific incompatibilities in the specific context of admixed populations, the *r*|*a* LD measure of positive ancestry disequilibrium (AD) was proposed (22). This measure is equivalent to the *r^2^_s_* in (18), and therefore corresponds to the *r^2^* measure corrected for the population structure (the *a* parameter being the genome-wide admixture proportion between two species). Although *r*|*a* reduced the false-positive rate in AD scans, an excess of low p-values was still observed in simulations of populations with different demographic histories with population structure (22).

In this study, we propose to implement a statistical test to detect epistatic selection in heterogeneously structured populations (i) between two bi-allelic SNPs by using the *r_v_* measure, or (ii) between two genomic regions including each multiple SNPs, by using the *cor_PC1v_* measure. This latter measure captures the quantitative correlation among the first principal components (PC1) summarizing the multi-locus genotypes at each genomic region. Compared with *r* and *cor_PC1_*, using simulations of genome-wide SNP data in structured diploid populations with random to self-mating schemes under two epistatic selection models, we showed that *r_v_* and *cor_PC1v_* (i) drastically reduce the background LD generated by population structure and relatedness among individuals; (ii) show an equivalent or a lower power to detect epistatic selection, depending on the mating scheme, on the dominance/codominance/recessivity of selected mutations and on whether coadapted or compensatory epistatic selection occurred; and (iii) *T*_*rv*_ or *T*_*corPC*1*v*_ statistics follow approximately a Student distribution τ_(*n*−2)_ under the null hypothesis of independence between the two tested loci. Hence, unlike *T*_*r*_ or *T*_*corPC*1_, *T*_*rv*_ or *T*_*corPC*1*v*_ can be used for straightforward statistical testing of the correlation coefficient between two loci, while accounting for spurious correlations due to population structure and heterogeneous relatedness among individuals.

Empirical detections of epistatic selection by genome-wide or targeted approaches on real SNP data are scarce in the literature (23–28). We assayed our new statistical test in the frame of genome-wide epistatic selection scans (GWESS) using a « bait» approach, with genomic SNP data from two different organisms: the model legume plant *Medicago truncatula* and human. As a proof of concept, we first described in *M. truncatula* the detection of epistatic selection between the Super Numeric Nodule *MtSUNN* gene, encoding a receptor which is central for the negative regulation of symbiotic root nodulation, and the CLAVATA3-like (CLE) signaling peptide *MtCLE02*. This finding is reminiscent of the functional interaction proposed for other CLE signalling peptides, *MtCLE12* and *MtCLE13* previously shown to require the *MtSUNN* receptor to regulate negatively the number of nodule organs on the plant root system (29, 30). Accordingly, an ectopic expression of the *MtCLE02* gene in *M. truncatula* wild-type and *sunn* mutant roots experimentally demonstrated a *MtSUNN*-dependent negative role of the *MtCLE02* gene on nodulation, hence validating functionally the genetic interaction between these two genes. In human, we illustrate the usefulness of the approach by identifying a significant epistatic or coselection signal in Central South Asian populations between *SLC24A5* and *EDAR* genes, encoding respectively a cation exchanger affecting pigmentation in zebrafish and human (31) and a receptor involved in the development of hair follicles, teeth and sweat glands (32, 33). Together with the fact that *SLC24A5* and *EDAR* were previously shown to be under strong positive selection in Europe and East Asian populations, respectively (34–36), our results highlight the role of epistatic selection or coselection in shaping gene coadaptation during the evolution of populations.

## Results

### Quality control of simulations

Details of our simulation procedure are provided in Materials and Methods. Briefly, genome-wide (4 chromosomes) SNP data (∼ 15,000 SNPs per chromosome) in a two-population split model with 250 diploid individuals per population during 300 generations (the ancestral population before the split was generated by coalescent simulations) were replicated 1,000 times for all combinations of the following parameter settings: (i) selection regimes as neutrality (NEUT), coadapted (COAD) and compensatory (COMP) two-locus epistatic selection, or additive (ADD) two-locus selection (all selection models starting 100 generations after the split time), (ii) random or self-mating (95% selfing rate) since the initial generation, and (iii) complete recessivity, codominance or dominance of the mutations under selection.

To ensure that our simulations produced consistent population structure and fixation of coselected alleles, we tracked down allele frequencies evolution at multiple SNP positions, including those of the derived alleles *a* and *b* at the two SNPs A and B intended to be targeted by selection, on two different chromosomes. Population structure and within population inbreeding were calculated using *F_ST_* and *F_IS_* parameters (**S1 Fig**). At the outcome of the simulations (i.e. generation 300), the average *F_IS_* in self-mating and panmictic populations was equal to 0.92 and 0.07 respectively, while the average *F_ST_* was equal to 0.19 and 0.10 respectively, thus showing realistic values. Selection efficiency was measured by the co-fixation rate of *a* and *b* at each generation in each selection model (**S2 Fig**). The first observation was that the COAD epistatic model generally induced a higher speed of fixation than the ADD positive selection model, while, as expected, the COMP model tends to maintain higher polymorphism due to selection of both *AB* and *ab* combinations at the two selected loci. The second observation was that rates of co-fixation of the derived alleles in self-mating populations reached more rapidly an equilibrium value than in panmictic populations, but more importantly that in self-mating populations the dominance, codominance or recessivity of the selected mutations had few effect on the co-fixation dynamics because of the very low heterozygosity level (*F_IS_* = 0.92 at the onset of selection). On the other hand, dominance, codominance or recessivity in panmictic populations strongly impact co-fixation dynamics due to more complex fitness patterns in the presence of heterozygotes (**S1 Table**). However, values of the fixation rates must be interpreted in light of the moderate sizes of the simulated population (N = 250 in each population), as selection efficiency increases with population size according to a factor *Ns* (37).

### Two-locus linkage disequilibrium under epistatic selection models

We focus hereafter on the evolution of the two-locus average LD across simulations, in self-mating and panmictic populations under selection models with **codominance**. We selected this genetic model because it produced the highest rates of co-fixation (*i.e.* the best selection efficiency) in panmictic populations, while similar co-fixation rates were observed under dominance, codominance or recessivity of the selected mutations in self-mating populations (**S2 Fig**). In addition, to perform proper comparisons between selection models and to avoid sampling bias on the average LD, in the COAD and ADD models we selected 500 simulations among those in which the outcome at the last generation was the co-fixation of the derived alleles *a* and *b* in both sub-populations, and in the COMP selection model we randomly sampled 500 simulations (*i.e.* simulations showing fixation of the *AB* or *ab* combination or still showing polymorphism at the last generation). For better visibility, the evolution of the average two-locus LD is described by using the absolute correlation values. Simulations under selection models with recessivity or dominance are provided in **S3 Fig**. In the codominant mutations model, we first observe that under the NEUT model population structure with or without non-random mating generates LD between two independent loci, as measured using *r* or *cor_PC1_*, which values can reach 0.25 to 0.5 at the final generation (**Fig 1**). This background LD is lowered to zero or close to zero on average, when correcting these statistics by the *V* matrix, as measured using *r_v_* or *cor_PC1v_*. This result makes the *r_v_* and *cor_PC1v_* of necessary use in order to perform statistical tests of the neutral hypothesis of a null correlation between two independent loci, accounting for “noisy” neutral processes (see next result section).

**Figure 1.**
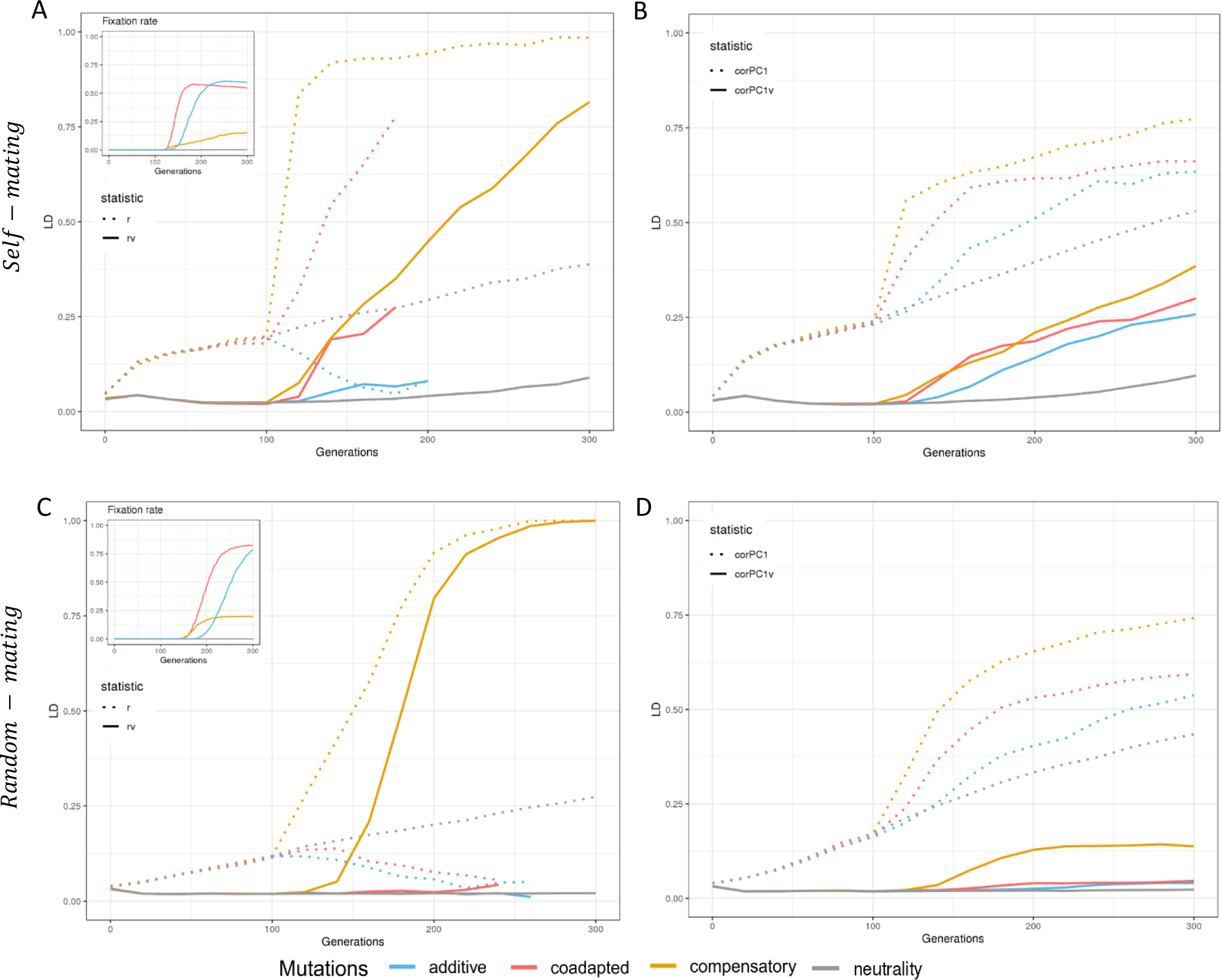
Evolution of inter-locus epistatic selection detected with linkage disequilibrium on simulated data. Evolution of inter-locus LD in self-mating simulation model calculated on SNP-based scale with *r* (*r_v_*) (**A**) and on window-based scale with *cor_PC1_* (*cor_PC1v_*) (**B**). Evolution of inter-locus LD in random mating simulation model calculated on SNP-based scale (**C**) and on window-based scale (**D**). Fixation rate in (A) and (C) depicted co-fixation of coselected mutant alleles *a* and *b* over generations, showing selection efficiency. Note that mutations under selection are codominant.

Second, we observe that selection models tend to generate more LD than NEUT model, though it depended on the statistics (*r*/*r_v_* or *cor_PC1_*/*cor_PC1v_*), the mating system and the selection model in an entangled way. For instance, the COAD, COMP and ADD selection models all tend to generate more LD than the NEUT model in self-mating species, with COAD and COMP generating more LD than the ADD model (**Fig 1**). In panmictic populations, however, only the COMP model generate consistent LD as measured using *r_v_* or *cor_PC1v_*, compared to the COAD and ADD models which generate few LD. A noticeable case is the one of LD measured in the ADD model with the *r* statistics, which is lower than in the NEUT model. This can be explained by a strong structure effect under the NEUT model, that is mitigated in the ADD model due to fixation independency of the derived alleles at the SNPs under selection. However, one can observe that it is not the case with LD measured by *cor_PC1_* which shows higher values than the NEUT model along the generations. This can be explained by a strong haplotype structure in the region surrounding SNPs under selection, between the two subpopulations, since haplotypes diverged during 100 generations before being targeted by selection. However, despite correcting for population structure, it remains difficult to distinguish epistatic selection from additive selection in self-mating populations at the haplotype level when these are not the same haplotypes that are under selection in the different subpopulations. Nonetheless, this nuisance phenomenon has less impact when focusing on the SNPs targeted by selection.

Finally, it can be seen that SNP-based LD measures (*r*/*r_v_*) are more efficient than haplotype-based LD measures (*cor_PC1_*/*cor_PC1v_*) to detect epistatic selection (**Fig 1**). However, they cannot capture any signal once allele fixation at one SNP or co-fixation at the two SNPs has occurred. On the other hand, *cor_PC1_*/*cor_PC1v_* rely on SNP polymorphism in the genomic region surrounding SNPs under selection, so that they can benefit from the hitch-hiking effect even after allele fixation at the SNPs targeted by selection.

### False positive control and detection power of two-locus correlation statistics

On the assumption that values of the correlation coefficient *r* follow a Student distribution τ_(*n*−2)_ under the null hypothesis of independence between the two variables tested, we examined the fit of the statistics *T*_*r*_, *T*_*corPC*1_, *T*_*rv*_ and *T*_*corPC*1*v*_ to such a distribution. This allowed evaluating their use for statistical testing of the null hypothesis that *T* = 0 – that is *r* = 0 – if the two tested loci are independent and evolve neutrally (see Materials and Methods for the formulas of the *T* statistics). False positive (FP) proportions of the statistics *T*_*r*_, *T*_*corPC*1_, *T*_*rv*_ and *T*_*corPC*1*v*_ with regard to the Student distribution τ_(*n*−2)_ are given for different rejection quantiles in **Table 1**.

**Table 1.** False positive (FP) proportions for *T*_*r*_, *T*_*corPC*1_, *T*_*rv*_ and *T*_*corPC*1*v*_ statistics in comparisons with the Student distribution (τ_(*n*−2)_) used for testing the significance of the correlation coefficient.

| Generation | Mating scheme | Statistics | FP proportions |  |  |
| --- | --- | --- | --- | --- | --- |
|  |  |  | 10% (1.283) | 5% (1.648) | 1% (2.334) |
| 140 | self-mating | $T_r$ | 85 % | 82 % | 74 % |
| | | $T_{corPC1}$ | 89 % | 86 % | 81 % |
| | | $T_{r_v}$ | 13 % | 8 % | 3 % |
| | | $T_{corPC1_v}$ | 13 % | 7 % | 3 % |
| | random-mating | $T_r$ | 72 % | 66 % | 55 % |
| | | $T_{corPC1}$ | 83 % | 78 % | 70 % |
| | | $T_{r_v}$ | 2.8 % | 0.6 % | 0.2 % |
| | | $T_{corPC1_v}$ | 2.5 % | 0.6 % | 0.1 % |
| 300 | self-mating | $T_r$ | 92 % | 91 % | 87 % |
| | | $T_{corPC1}$ | 95 % | 93 % | 91 % |
| | | $T_{r_v}$ | 31 % | 26 % | 20 % |
| | | $T_{corPC1_v}$ | 37 % | 31 % | 22 % |
| | random-mating | $T_r$ | 85 % | 81 % | 74 % |
| | | $T_{corPC1}$ | 93 % | 91 % | 87 % |
| | | $T_{r_v}$ | 4.6 % | 2.3 % | 1.2 % |
| | | $T_{corPC1_v}$ | 5.2 % | 3.1 % | 1.1 % |

False positive proportions are calculated as the proportion of simulations in which the statistics has a value greater than the defined rejection quantile of the $\tau_{(n-2)}$ distribution, for different type I errors: 10%, 5% and 1% (the associated rejection quantile are given in bracket). In our simulations, the sample size $n$ was equal to 500. Since the sign of the correlation coefficient is not interpretable, especially for $T_{corPC1}$ and $T_{corPC1_v}$ the absolute values of $T_r$ , $T_{corPC1}$ , $T_{r_v}$ and $T_{corPC1_v}$ and of the Student distribution $\tau_{(n-2)}$ were used for false positive proportion calculation.

Two time points were considered in the neutral simulations for the false positive analysis as well as power analysis, at generation 140 in the midst of the time course and at the last generation 300. At generation 140, in both the self-mating and random-mating models, *T*_*r*_ and *T*_*corPC*1_ showed excessively large FP proportions. For instance, FP proportions ranging from 55 to 81% were observed for a 1% type I error, while *T*_*rv*_ and *T*_*corPC*1*v*_ showed adequate, conservative FP proportions ranging from 0.1 to 3% for the same 1% type I error (**Table 1**). At generation 300, a similar behaviour was observed, with FP proportions ranging from 74 to 91% for a 1% type I error, while *T*_*rv*_ and *T*_*corPC*1*v*_ showed adequate – though less conservative in the case of the self-mating model – FP proportions ranging from 1.1 to 22% for the same 1% type I error. This indicates that correction for population structure and heterogeneous relatedness is crucial for correctly testing the null hypothesis of the correlation coefficient between two SNP or two genomic regions. However, while sharp population structure is efficiently taken into account in the statistical correction, it seems that the combination of such population structure and self-mating tend to inflate the distribution of *T*_*rv*_ and *T*_*corPC*1*v*_ on the long term, since this phenomenon was observed only at generation 300 and not at generation 140.

Then, a power analysis was performed for the statistics *T*_*r*_, *T*_*corPC*1_, *T*_*rv*_ and *T*_*corPC*1*v*_ by using (i) simulated data under the null hypothesis of neutrality and independence between loci and (ii) simulated data under each of the selection model and independence between loci. For each statistic, neutral simulations were used to estimate one-way rejection quantiles by using the absolute values of the statistic, corresponding to type I error α ranging from 0.001 to 0.20. Then, for each selection model under self-mating or random mating with codominant mutations, we calculated the proportion of simulations where absolute values of each *T* statistic were higher than the rejection quantiles. Power was calculated for *T*_*r*_, *T*_*corPC*1_, *T*_*rv*_ and *T*_*corPC*1*v*_ at generation 140 where allele fixation at SNPs under selection was not yet achieved, and also at the last generation 300 for window-based measures *T*_*corPC*1_ and *T*_*corPC*1*v*_.

At both time-point, a general trend is that the detection power with *r*/*r_v_* and *cor_PC1_*/*cor_PC1v_* is higher for the COMP model than for the COAD or the ADD models (i.e. 25-50%, 10-65%, and 10-30%, respectively, for α=5% with *r_v_* or *cor_PC1v_* statistics), especially when considering random mating (**Fig 2**). In addition, the correction of LD-based measures by the kinship matrix (*r_v_*/*cor_PC1v_*) does not increase detection power of epistatic selection; rather it tends to reduce power especially in the COMP model but not in the COAD model. This is due to the fact that the fixation of the *AB* allelic combination is more frequent in sub-populations than in the whole population in the COMP model (see **S2 Fig**) – a consequence of unequal initial frequencies of the ancestral/derived alleles in the simulations – leading to a high LD value when population structure is not taken into account. On the other hand, it is interesting to note that an increased power to detect additive selection is observed with *r_v_* especially in a self-mating regime (**Fig 2A**). This is because, at the whole population level, the three allelic combinations containing the derived allele (*i.e. ab*, *Ab* and *aB*) are selected, while in sub-populations the self-mating regime limits this selection to the most frequent combinations that are present at the onset of selection, thus increasing LD values when population structure is taken into account. Finally, *cor_PC1v_* may tend to show less power than *r_v_* when the same allele selected in each subpopulation is present on different haplotypes, which is the case in our simulations (**Fig 2A,C; Fig 2B,D**, and see previous section).

**Figure 2.**
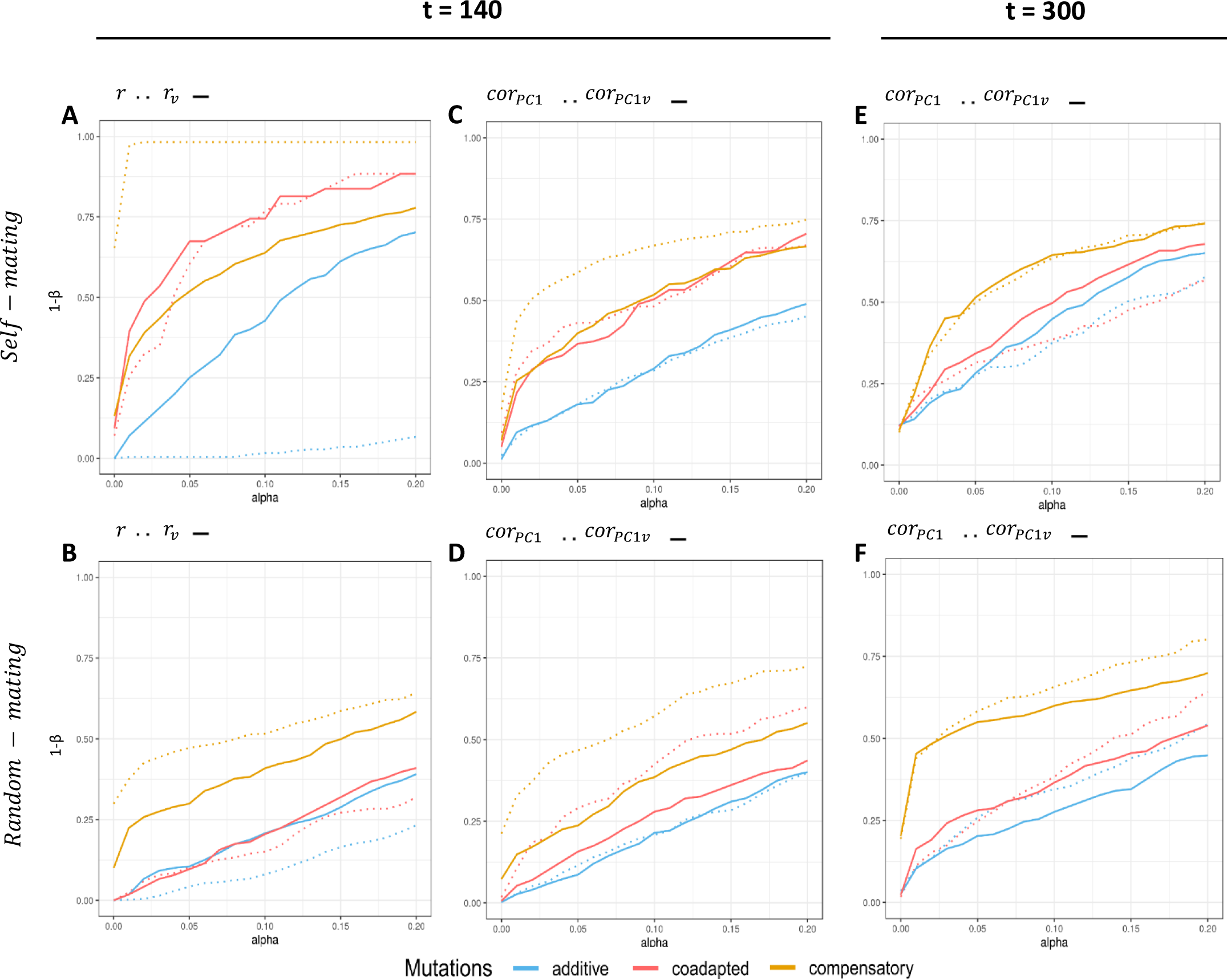
Detection power of epistatic selection according to neutral models for SNP-based and window-based LD measures. Detection power of epistatic selection in self-mating simulation model and in random mating model calculated on SNP-based scale (*r* and *r_v_*) (**A, B**) and on window based-scale (*cor_PC1_* and *cor_PC1v_*) (**C**, **D, E, F**). Figures (**A, B, C, D**) depict the detection power at generation 140 and figures (**E**, **D**) at generation 300 (*r* and *r_v_* are no longer computable at this generation in coadapted and additive selection models). The *x*-axis corresponds to the type I error (α) and y-axis corresponds to the detection power (1-β). Mutations under selection are codominant.

### Detection of two-locus coadaptation in the plant *Medicago truncatula*

To illustrate the straightforward statistical testing of the correlation coefficient between two loci using *T*_*corPC*1_ or *T*_*corPC*1*v*_, a one-dimension Genome-Wide Epistatic Selection Scan (GWESS) was performed using a bait approach with the *MtSUNN* candidate gene against 48,339 other genes of the *M. truncatula* genome. The statistics *T*_*corPC*1_ and *T*_*corPC*1*v*_ were calculated based on PC1 values from SNP data located in 10 kbp windows spanning each *M. truncatula* gene and PC1 values from *MtSUNN*, and p-values were obtained from the τ_(*n*−2)_ null distribution. Two scans were implemented, including SNP data from either the whole *M. truncatula* collection – n=262 individuals – or from the Far West (FW) sub-population – n=80 individuals – (**Fig 3A,C; Fig 3B,D**; respectively). A clear inflation towards small p-values can be observed for scans based on *T*_*corPC*1_ (**Fig 3A,B**) compared with scans based on *T*_*corPC*1*v*_ (**Fig 3C,D**), and this inflation is higher with data from the whole collection which shows a higher degree of population structure. In the FW sub-population scan, a sharp peak is observed using *T*_*corPC*1*v*_ on the chromosome 6 corresponding to the *MtCLE02* (Medtr6g009390) gene on top of the peak (**Fig 3D**, p-value = 1.7×10^−8^). *MtCLE02* corresponds to the top candidate gene showing an epistatic selection signal outside of the chromosome 4 where is located *MtSUNN*. Whereas *MtCLE02* is also highly correlated with *MtSUNN* when considering *T*_*corPC*1_ (**Fig 3B**, p-value = 2.74×10^−13^), several other genomic regions display similar or even more significant signals, which may indicate spurious genome-wide correlations. Interestingly, scans based on SNP data from the whole *M. truncatula* population do not reveal such strong signal in the genomic region containing *MtCLE02* (**Fig 3A,C;** p-value = 0.077 and 0.006 for *T*_*corPC*1_ and *T*_*corPC*1*v*_, respectively), indicating that in this specific case, epistatic selection may have occured at the sub-population level, and as such could not be detected using SNP data from the whole population and the *T*_*corPC*1*v*_ statistic because the signal would be correlated with population structure.

**Figure 3.**
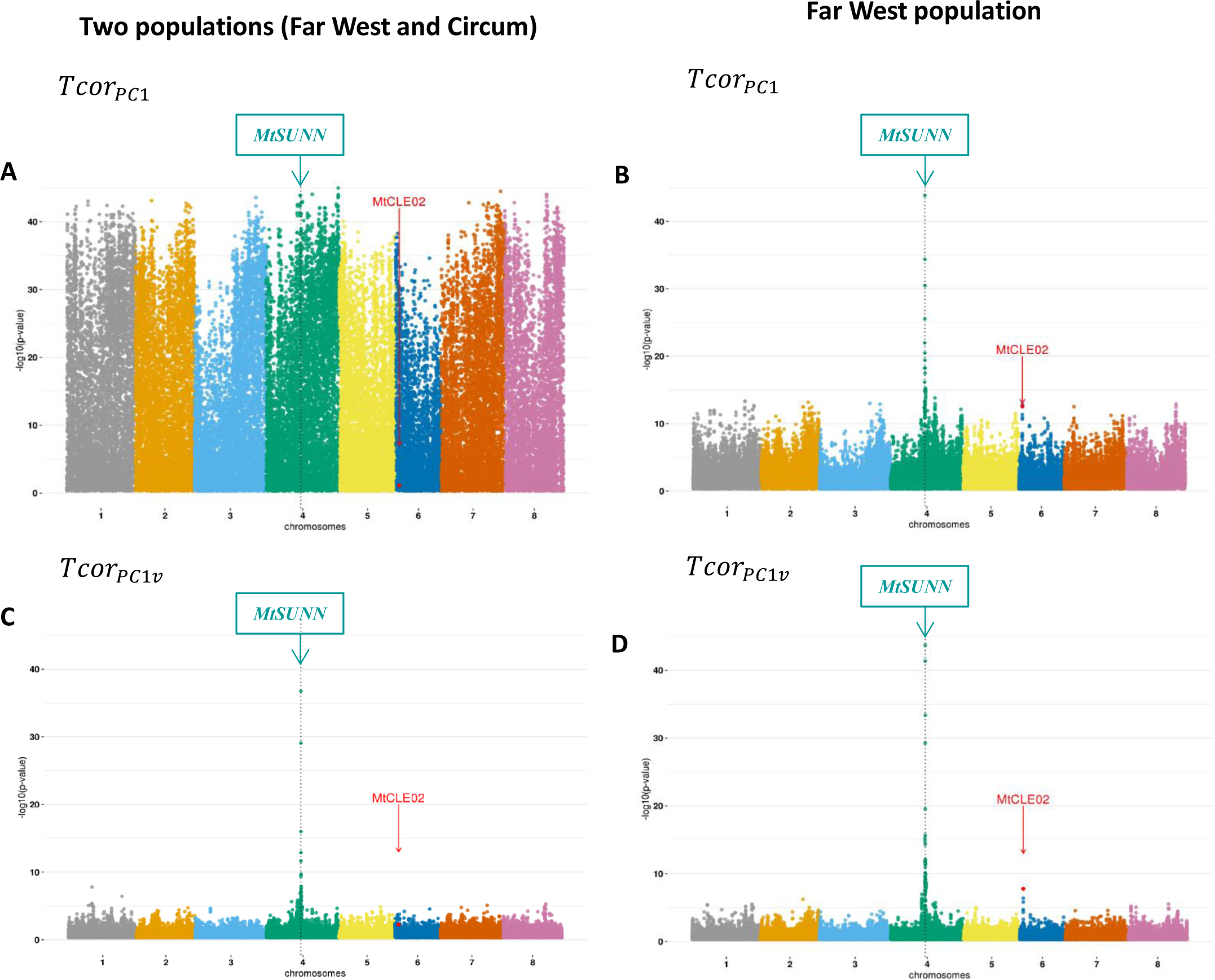
LD distribution between the bait gene *MtSUNN* and all genes of *M. truncatula* genome. LD between *MtSUNN* gene (framed) and all *M. truncatula* genes was calculated in the entire population (**A** – **C**) and in the Far-West population (**B** – **D**). The p-values of the correlation test were calculated from *Tcor_PC1_* correlation (**A** – **B**) and from *Tcor_PC1v_* statistics **(C** – **D**), which includes the kinship matrix. The *x*-axis corresponds to genes positions spanning the 8 chromosomes, each point corresponding to a gene and red dots depicting the *MtCLE02* gene in each figure. The y-axis shows the −log_10_(p-value) of the test of the correlation coefficient.

### Experimental evidence for the genetic dependency between *MtSUNN* and *MtCLE02* in *Medicago truncatula*

The *MtSUNN* gene encodes a Leucine-Rich Repeats – Receptor Like Kinase (LRR-RLK) whereas the *MtCLE02* genes encodes a CLAVATA-like secreted signalling peptide. The SUNN receptor function, which is crucial in the systemic negative regulation of nodulation, has been previously associated to other CLE secreted signalling peptide encoding genes, *MtCLE12* and *MtCLE13* (29, 38). Whereas their expression was induced by the inoculation with symbiotic rhizobia bacteria initiating nodulation, this was not the case for *MtCLE02* (**S4A,B Fig**). These previously documented CLE/SUNN relationships pointed us to test for a putative functional interaction between CLE02 signalling peptides and the SUNN receptor. As previously performed for *MtCLE12* or *MtCLE13* (29, 38), we used a genetic approach consisting in overexpressing the *MtCLE02* gene comparatively in *M. truncatula* Wild-Type (WT) and *sunn* mutant roots. First, quantification of the nodule number in WT versus *sunn* mutant roots highlighted the well-known supernodulation phenotype of the *sunn* mutant (Mann & Whitney – Wilcoxon test, p-value = 2×10^−6^). Second, the nodule number was significantly decreased when *MtCLE02* was overexpressed in WT roots, as validated by real time RT-PCR (**S4C Fig;** Mann & Whitney – Wilcoxon test, p-value = 2×10^−6^), indicating a negative role of *MtCLE02* on nodulation (**Fig 4A,B)**. Third, *MtCLE02* overexpression in *sunn* mutant roots did not impact the nodule number (Mann & Whitney – Wilcoxon test, p-value = 0.66), in contrast to what was observed in the WT, indicating that the negative role of CLE02 on nodulation relies on the SUNN receptor. These experimental results therefore unambiguously functionally link these two *M. truncatula* genes in the context of symbiotic root nodulation through an epistatic relationship which was detected by our method based on their putative co-evolution.

**Figure 4.**
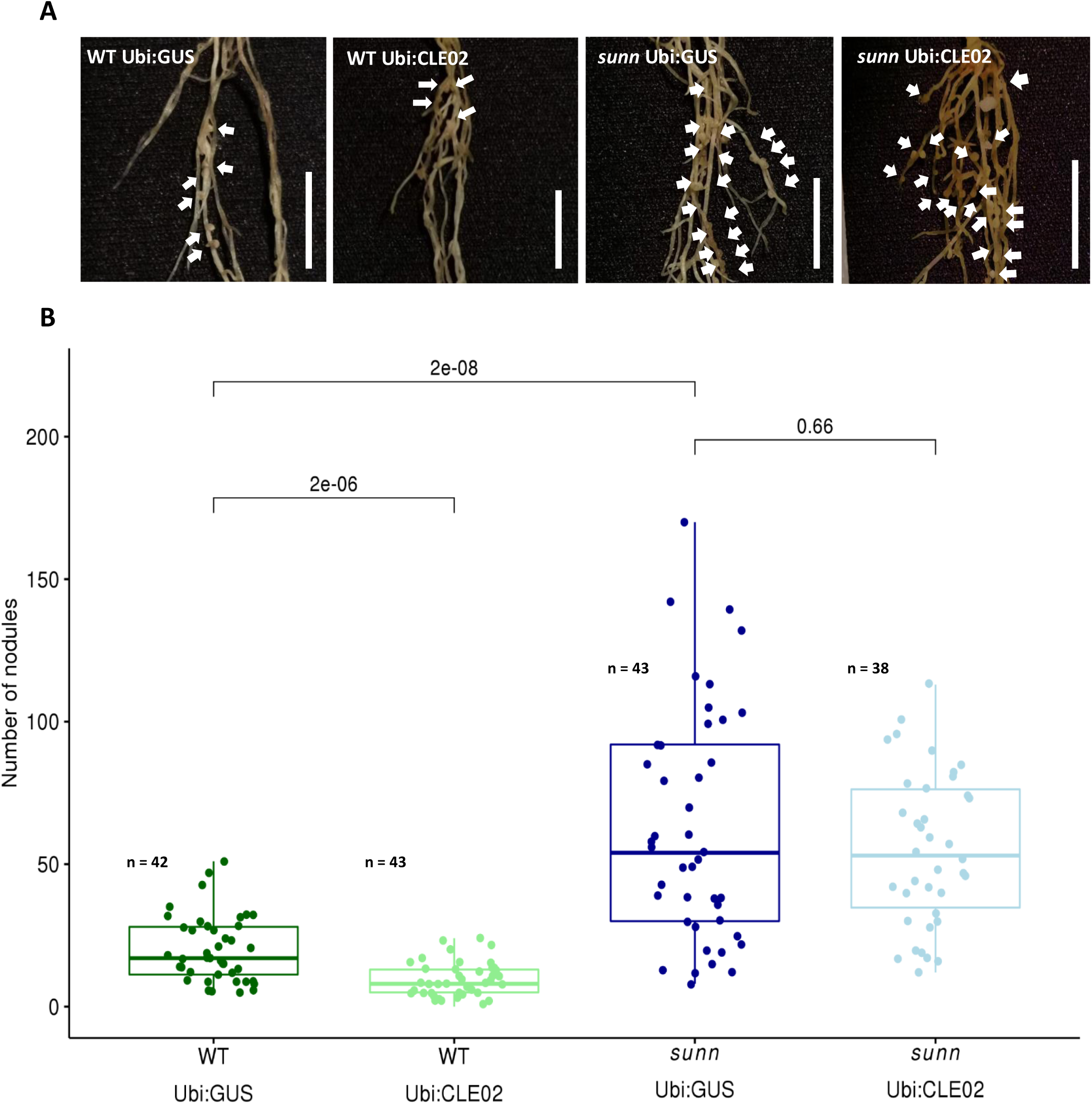
Experimental validation of the MtCLE02 signalling peptide / MtSUNN receptor functional relationship in *M. truncatula* symbiotic nodulation. **(A)** Representative images of nodulated roots, 14 days post rhizobium inoculation, overexpressing the *MtCLE02* gene (Ubi:CLE02) or a *GUS* control gene (Ubi:GUS) either in Wild-Type (WT) plants or in the *sunn* mutant. Scale bar = 1 cm. **(B)** Boxplots of the number of nodules in the same conditions as described in **A**. A Mann & Whitney Wilcoxon rank sum test was used to assess pairwise statistical differences, as indicated within the graph.

### Detection of two-locus coadaptation in human populations

In human, two GWESS were performed on the world-wide sample of 940 individuals with two bait SNPs, 15_46172199 (rs2250072) and 2_108973688 (rs6749207) located in *SLC24A5* and *EDAR* genes, respectively. For each bait SNP, SNP-based statistics *T*_*r*_ and *T*_*rv*_ were calculated for 479,764 genome-wide SNPs. Scans implemented in the world-wide population with both 15_46172199 and 2_108973688 SNPs as bait were inflated towards small p-values when using *T*_*r*_ statistic (**Fig 5A,B**), compared with scans implemented with *T*_*rv*_ statistic (**Fig 5C,D**). Using SNP 15_46172199 as bait for *SLC24A5* gene, a peak corresponding to the *EDAR* gene was detected, with SNP 2_108946170 as the top significant SNP (**Fig 5A,C**; *T*_*r*_-based p-value = 2.29×10^−9^). Conversely, when the scan was performed with SNP 2_108973688 as bait from the *EDAR* gene, a peak corresponding to the *SCL24A5* gene was detected, with SNP 15_46179457 as the top significant SNP (**Fig 5B,D**; *T*_*rv*_-based p-value = 1.2×10^−12^). Genome-wide LD distributions between each bait SNPs and all other SNPs (**Fig 5**; top left of each panel) showed that LD values between SNPs from *EDAR* and *SLC24A5* locate in the extreme right tail thus indicating extremely significant signals. The world-wide geographical distribution of genotypes at SNPs 15_46172199 – *SLC24A5* – and 2_108973688 – *EDAR* – (**Fig 6C**) correlates substantially with global population structure, as depicted by a phylogenetic tree based on the kinship matrix among individuals (**Fig 6A,B**). Indeed, the derived allele at SNP 15_46172199, associated with the light skin allele at the *SCL25A5* gene is present in Europe, North Africa, Middle East and Central South Asia, and the derived allele at SNP 2_108973688, associated with the thick hair allele at the *EDAR* gene is present in East-Asia, America, and Oceania (**Fig 6C**). The strong LD signature between *SCL25A5* and *EDAR* observed in the world-wide sample, as measured with *T*_*r*_, therefore reflects selection of derived alleles (in green) in different geographical regions (34, 36), thus a correlation with global population structure. However, LD corrected for population structure and relatedness between individuals, as measured with *T*_*rv*_, was still highly significant between *SCL25A5* and *EDAR*, indicating that coadaptation by coselection or epistatic selection may have occurred between both genes at the level of geographical sub-regions. In order to localize the geographic origin of such selection signatures, GWESSs were performed within six geographical regions of the world-wide sample: Central South Asia, East Asia, Subsaharian Africa, Middle East, Europe and America (**S5-S10 Figs**). Only the GWESS performed in Central South Asia indicated significant LD between SNPs at the *SCL24A5* and *EDAR* genes (**S5C Fig**, *T*_*rv*_-based p-value = 6.7×10^−6^ at SNP 2_108973688; **S5D Fig**, *T*_*rv*_-based p-value = 2.8×10^−6^ at SNP 15_46174380). Human population samples from the HGDP-CEPH dataset in Central South Asia are composed of eight different ethnies from Pakistan. Scans implemented with *T*_*r*_ or *T*_*rv*_ statistics showed similar genome-wide LD distributions (**S5 Fig**; top left of each panel), indicating a weak population structure within this geographic region. To search for local signals of coselection or epistatic selection, LD tests were performed with *T*_*rv*_ between two candidates SNPs within *SCL24A5* (15_46179457 (rs1834640) and 15_46172199) and three candidates SNPs within *EDAR* (2_108962124 (rs260607), 2_108982808 (rs17034770) and 2_108973688), for the 50 ethnic groups or populations distributed in the eight geographical regions and showing polymorphism at all five SNPs. Average and standard deviation of –log_10_(p-value) across six pairwise SNP comparisons (**Fig 6,D**) strongly support high LD between *SCL24A5* and *EDAR* in the Burusho ethnic group from Pakistan (3.2 and 0.36, respectively), as also highlighted by genotypes of Burusho individuals (**Fig 6,C**, *r*_*v*_ = 0.63 for genotypes between the two dotted lines). This pattern of high LD between *SCL24A5* and *EDAR* in Burusho seems not to be generated by population sub-structure in this ethnic group since LD tests performed with *T*_*r*_ result in similar average (3.18) and standard deviation (0.36) of –log_10_(p-value) (**S11 Fig**). These analyses also suggest a weaker but still significant signal in the Hazara ethnic group, with average and standard deviation of –log_10_(p-value) of 1.73 and 0.14, respectively.

**Figure 5.**
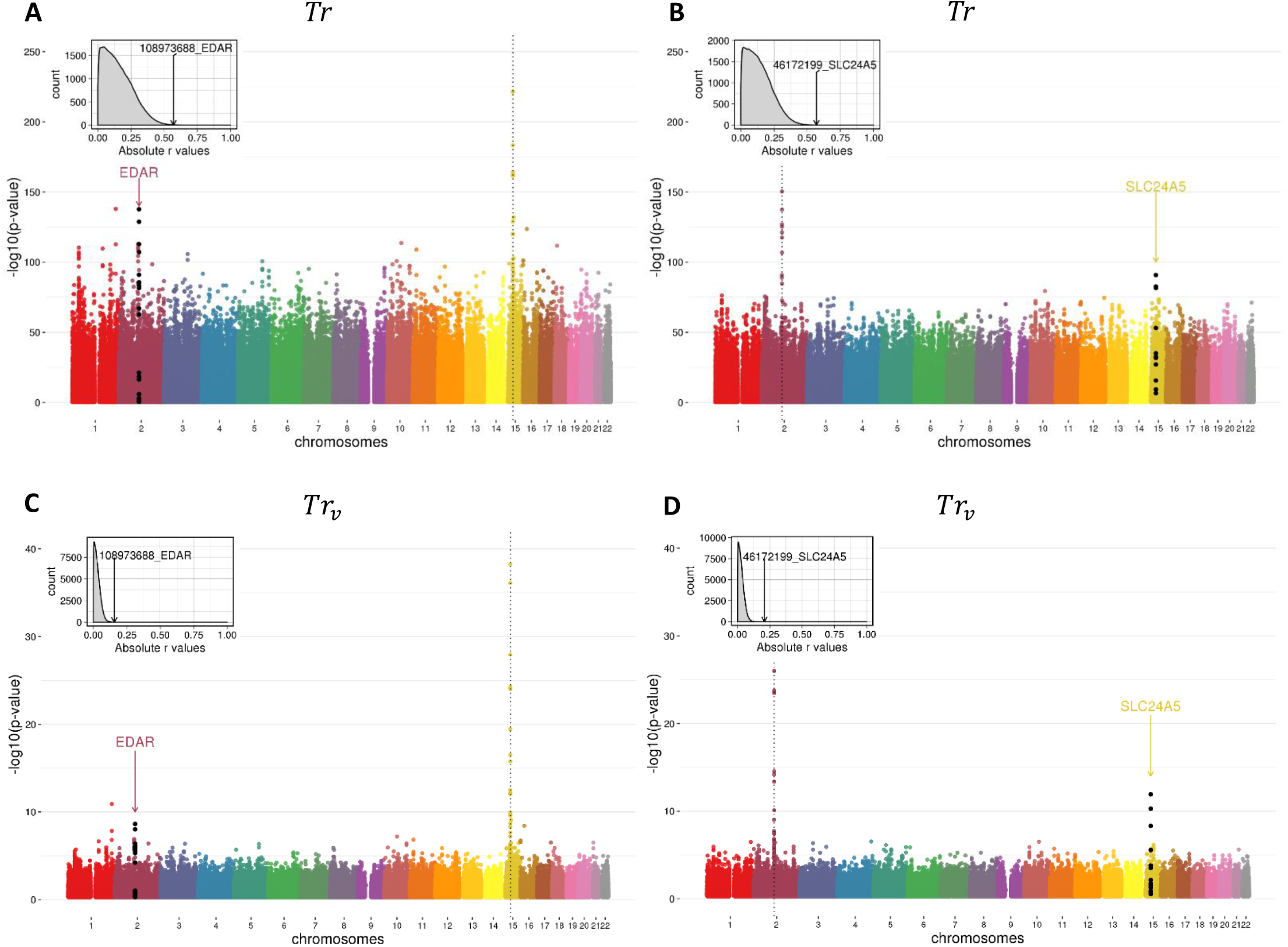
LD distribution between the bait SNPs of *SLC24A5* and *EDAR* genes and all other HGDP-CEPH SNPs in the whole human population samples (n=952). LD between SNP_15_46172199 (*SLC24A5)* and SNP_2_108973688 (*EDAR*), respectively, and all other SNPs of the genome is tested using *T*_*r*_ (**A, B**) or *T*_*rv*_ (**C, D**), which includes the kinship matrix. The *x*-axis corresponds to SNP positions spanning the 22 human autosomes, each point corresponds to a SNP and the black points depict SNPs at candidate genes in epistatic selection with one SNP at the bait gene (vertical dotted line) in each figure. The y-axis is the −log_10_(p-value) of the test of the correlation coefficient. Plots at the top left of each figure show distribution of LD between each bait SNP and all other SNPs of the genome. LD value between bait SNPs of *SLC24A5* and the target SNP of *EDAR* (respectively bait SNPs of *EDAR* and target SNP of *SLC24A5*) is represented by an arrow.

**Figure 6.**
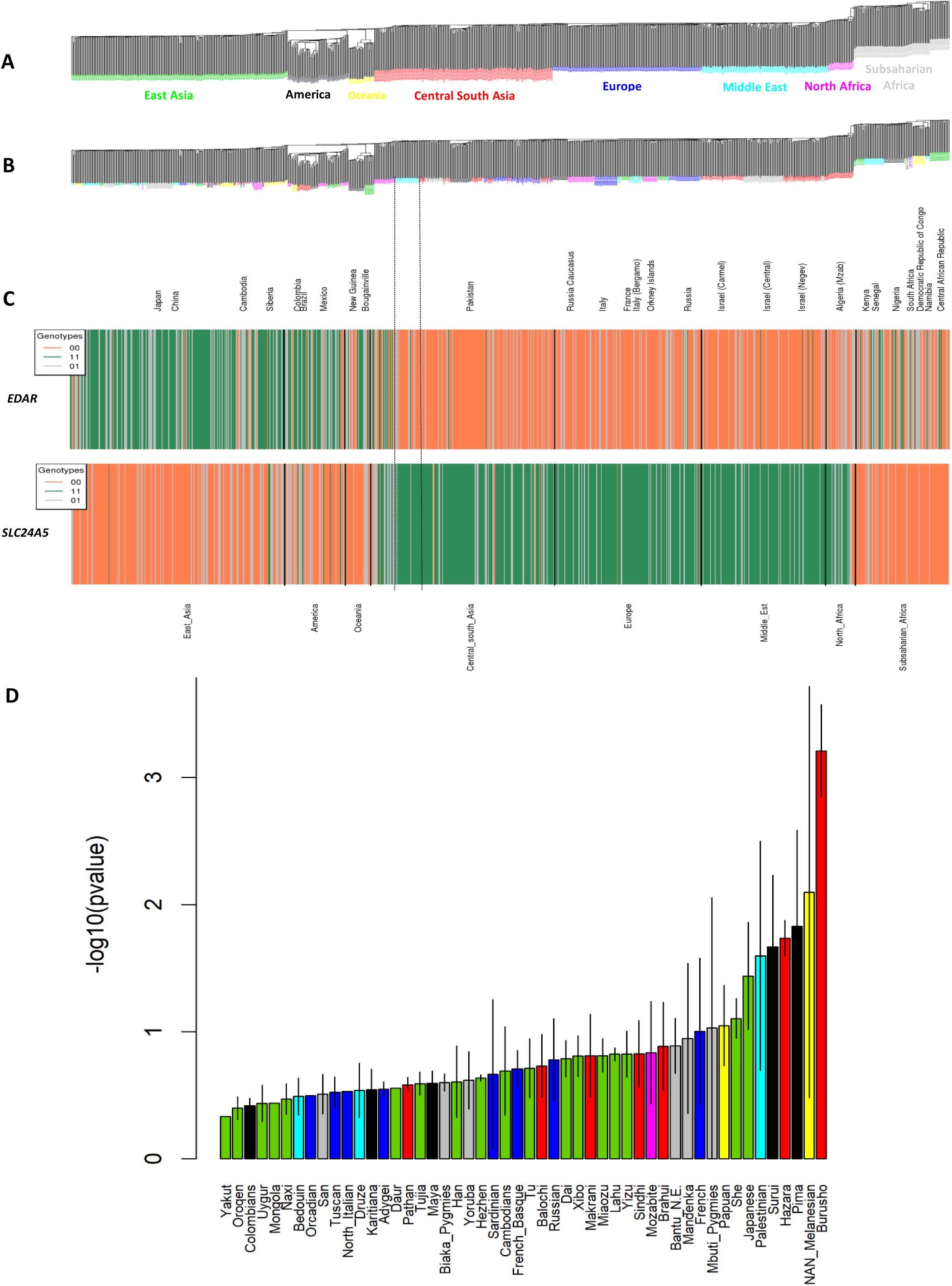
Schematic of human population structure inferred from kinship matrix, geographic distribution of alleles and LD between SNP 15_46172199 (*SLC24A5)* and SNP 2_108973688 (*EDAR*). (A) Neighbor-Joining tree inferred from the molecular kinship matrix based on 494,764 SNPs from HGDP-CEPH showing global human population structure. (B) Same tree as in (A) showing the clustering of the different sub-populations or ethnies sampled. (C) Barplots depicting the geographic distributions of genotypes at SNP 2_108973688 (*EDAR*) and SNP 15_46172199 (*SLC24A5*) highlighting the LD patterns mainly due to global population structure and selection of the derived allele (coded 1) at the two genes. (D) Average and standard error of LD significances based on *T*_*rv*_ statistics between SNPs of *SLC24A5* (SNP 15_46172199, SNP 15_46179457) and *EDAR* (SNP 2_108962124, SNP 2_108973688, SNP 2_108982808) within each human sub-population. The barplot pinpoints Central South Asia as the main source of within population LD probably due to coselection of alleles in Pakistan ethnies (mainly from Burusho, delimited by two vertical dotted lines in (C)).

## Discussion

We introduced a straightforward statistical method which can detect the signature of epistatic selection using LD between two loci, in order to identify genes that coevolve. SNP-based (*T*_*rv*_) and window-based (*T*_*corPC*1*v*_) statistics, which take into account the underlying population structure and relatedness among individuals, are shown to fit a Student distribution τ_(*n*−2)_, allowing to easily and quickly test for significance of correlation coefficients in the frame of Genome-Wide Epistatic Selection Scans (GWESS) using either a candidate SNP, a gene, or a short genomic region as bait. Simulations have shown that *T*_*rv*_ and *T*_*corPC*1*v*_ showed equivalent or less power than *T*_*r*_ or *T*_*corPC*1_ to detect epistatic selection in structured populations, ranging from 10 to 65% depending on the epistatic selection model and mating scheme (assuming a 5% type I error), and in the case where selection occurred simultaneously in all sub-populations. Thus, selection signals in local populations could be more difficult to detect with *T*_*rv*_ and *T*_*corPC*1*v*_ because in these cases selection can be correlated with population structure. In addition, the contribution of each sub-population to the global LD signal may not be easy to partition when heterogeneous relatedness and complex population structure underlie the genotypic data. Furthermore, the impact of the kinship matrix on LD correction changes depending on the scale of the sampling, with a stronger impact for large scales of geographical population structure than for smaller, less structured, and less heterogeneous geographical scales. This result therefore suggests to perform GWESS not only using a global sample comprising individuals from different populations, but also using samples from different sub-populations in order to search for more population-specific patterns of epistatic selection. In the context of statistical testing, however, simulations as well as analyses performed both in *M. truncatula* and human genomes strongly support the requirement of using *T*_*rv*_ and *T*_*corPC*1*v*_ instead of only *T*_*r*_ or *T*_*corPC*1_, in order to efficiently control for false positives in the frame of GWESS. Finally, despite SNP-based statistics (*T*_*r*_, *T*_*rv*_) may tend to show better detection power than window-based statistics (*T*_*corPC*1_, *T*_*corPC*1*v*_) because adaptive interacting mutations at both loci can be directly tested with SNP-based statistics and they are not sensitive to complex haplotype structure, window-based statistics also show several advantages. They are faster to implement at the genome scale – especially if two-dimensional GWESS are envisioned -, the window size can be fixed at a value that fits best the average LD decay in the species studied (even though a standard 10 kbp size can be used by default), and they allow to detect coevolving genes even after putative fixation of coselected SNPs due to surrounding SNPs within genes or windows that also carry a selection signal by hitchhiking.

The applications of our method to SNP data from human populations or from populations of the model plant *Medicago truncatula* allowed in both cases to identify couples of genes most probably under epistatic selection, or at least under coselection. In *M. truncatula*, polymorphism at the *MtSUNN* gene could be driven by a balancing selection at the local level because the *H* statistic (39) is 1.45 in the Far West population and ranks among the highest 8.41% in the entire genome, while *H* = 0.52 based on the whole species, ranking among the highest 39.5%, according to a set of 47,875 genes. The polymorphism at the *MtCLE02* gene seems however more affected by an ongoing selective sweep that can be detected at the level of the whole species (*H*= −2.25, ranking among the lowest 8.98% of the genome; data not shown). Still, both genes maintain polymorphism, and epistatic selection could drive this pattern probably through a compensatory model. As a proof of concept to illustrate the putative role of epistatic selection in the coadaptation of these two genes, a genetic approach was used to demonstrate the functional interaction between the MtCLE02 signalling peptide and the MtSUNN receptor. Indeed, the MtCLE02 signalling peptide negatively affects nodule number depending on the MtSUNN receptor, as previously shown for other CLE peptide encoding genes (*MtCLE12* and *MtCLE13*) (29, 38). Interestingly, it should be noted that the *MtCLE02* gene is, in contrast to *MtCLE12* and *MtCLE13*, not regulated by symbiotic nodulation conditions and not phylogenetically closely related to these previously characterized CLE signalling peptide encoding genes shown to have a related negative impact on nodule number (40). The identification of a novel CLE peptide acting in this genetic pathway highlights the discovery power of our method to functionally associate gene pairs independently of their expression pattern or of a coregulation pattern criterion.

In human, the *SLC24A5* gene, a major driver of variation in skin pigmentation, has been shown to be under positive selection in European population (36, 41, 42). The causal mutation for the light skin phenotype was not present in HGDP data (SNP rs1426654, position: 15_46213776), but the SNPs we used (i.e. 15_46179457 and 15_46172199) localize on the same *SLC24A5* haplotype background previously characterized (43–45). In addition, the V370A mutation in *EDAR* coding for a receptor related to TNFα receptors involved in driving hair structure and teeth and sweat glands development, was shown to be under positive selection in East Asia and in native Americans, and to increase hair thickness (33–35). Likewise, the causal mutation in the *EDAR* gene was not present in HGDP data, but the SNPs used (i.e. 2_108962124, 2_108973688 and 2_108982808) locate within the genomic sequence of *EDAR*. The geographic distribution of genotypes at SNP 15_46172199 and 2_108973688 strongly correlates with world-wide human population structure, which explains the high LD observed at this level in the scans implemented with *T*_*r*_. However, scans implemented with *T*_*rv*_ also indicate strong LD between *SLC24A5* and *EDAR* in the world-wide sample suggesting a persistence of LD due to epistatic selection or coselection. We identified Central South Asia, with the Burusho ethnic group from Pakistan as being the most probable geographic origin of such selection signature. GWESS performed in Central South Asia, and subsequent LD tests between *SLC24A5* and *EDAR* in sub-populations performed with *T*_*r*_ or *T*_*rv*_ statistics showed similar results, indicating weak population structure, as previously observed within this geographic region and in India (46). The Burusho ethnic group shows a predominant association between the derived alleles at *SLC24A5* and the ancestral alleles at *EDAR*, which is indicative of the persistence of a typical European light skin and thin hair structure in Burusho, coexisting with probably fewer East Asian morphotypes with darker skin and thicker hair structure. This pattern could have been driven by either epistatic selection, if a functional link between skin pigmentation and the ectodysplasin pathway occurred in human, or coselection for these two phenotypic traits, which is more likely because such a pattern seems specific from ethnic groups. Thus, in human, LD between *EDAR* and *SCL24A5* is not only related to world-wide population structure and positive selection but also to local coselection of genes.

A natural extension of the one-dimension use of *T*_*rv*_ or *T*_*corPC*1*v*_ – based tests in GWESS is the implementation of two-dimensional GWESS in which the correlation of each polymorphic locus (SNP, gene or genomic region) in the genome would be tested against all remaining polymorphic loci. Such type of analyses will open the way towards exploring the structure and topology of coadapted gene networks.

## Materials and Methods

### Genetic models of epistatic selection

We follow fitness genotype formalization under epistatic selection models as in (10, 13). Consider two independent bi-allelic loci A and B, with ancestral alleles *A* and *B*, and derived alleles *a* and *b*. The coadaptation model consists in positively selecting the two-locus *ab* combination; it can be considered as a selective sweep on the *ab* combination. The compensatory model consists in selecting against the *Ab* and *aB* two-locus combinations, but not against *AB* and *ab*. The coefficient *s* is used to positively or negatively select two-locus genotypes. In a diploid population, the two-locus fitness expression will depend on the level of dominance of the derived alleles (**S1 Table**). In the compensatory model with codominance, we choose to select against the double heterozygote genotype (*Aa*/*Bb*) because A and B are independent loci putatively equally expressing *AB* / *ab* or *Ab* / *aB*. In the neutral model, all fitness values are set up to 1.

### SNP-based and window-based LD measures of epistatic selection

In a diploid organism, at a given bi-allelic SNP with alleles coded 0 and 1 the three possible genotypes are (00,01,11), which can be coded as the allelic dose of allele 1 (0, 1, 2). The measure on unphased genotypes between two bi-allelic loci is defined by the correlation coefficient *r* between the vectors of genotypes at the SNPs *l* and *m*, *X^l^* and *X^m^* (47–49):

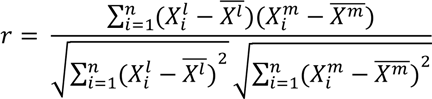

In the case where 0 and 1 are the ancestral and derived alleles, respectively, the sign of *r* indicates whether combinations of ancestral and derived alleles (i.e. 00 and 11) preferentially segregate in individuals at the two SNPs or whether alternative combinations do segregate preferentially (i.e. 01 and 10). At two physically unlinked loci in a panmictic population, this measure allows to detect fitness interaction between two new mutations under the coadaptation or the compensatory model (13). In the context of GWESS with high-density SNP data, we propose to use the *cor_PC1_* measure of LD between to genomic regions containing each multiple SNPs. The first principal component *PC1^l^* is used to summarize quantitatively the multi-SNP genotypes of the genomic region *l* (see (50)). Then, *cor_PC1_* is the correlation coefficient between the vectors of summarized multi-SNP genotypes of the two genomic regions *l* and *m*, *PC1^l^* and *PC1^m^*:

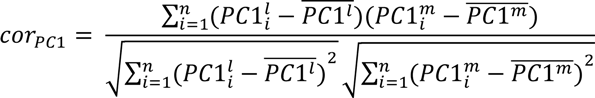

However, as mentioned in (18), population structure and relatedness among individuals generate non-independence between individuals and tend to bias upwardly the values of linkage disequilibrium measures. This is particularly the case in highly inbred or predominantly selfing species (20). At a given locus, Mangin and collaborators proposed to uncorrelate the observations by multiplying the vector of genotypes by 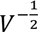, where *V* is the kinship (or relatedness) matrix among individuals. *V* is built with the *V*_*ij*_ covariance for all pairs (*i*,*j*) of individuals. *V*_*ij*_ is the average number of identical genotypes between individuals *i* and *j*, in a genome-wide SNP dataset. This measure of Identity In State (IIS) is a good proxy of Identity By Descent (IBS) with SNP markers since the latter follow an infinite site mutation model. It follows that, since *r* is the Pearson correlation coefficient, *r_v_* can be computed as 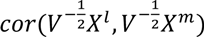 and *corPC1v* as 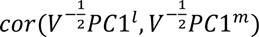.

### Statistical test of epistatic selection based on linkage disequilibrium

Under the hypothesis that observations within *X^l^* and *X^m^* (respectively within *PC1^l^* and *PC1^m^*) are independent, then *r* (respectively *cor_PC1_*) can be used to obtain the *T* statistics:

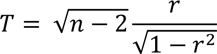

which follows a Student distribution τ_(*n*−2)_. However, in the case were observations are not independent; that is when the genotypes at a given locus are correlated within the population due to non-random mating and/or between populations due to structure, then we expect that only the *T* statistics obtained using *r_v_* or *cor_PC1v_* follow the distribution τ_(*n*−2)_. In the case where the ancestral/derived allele status is known at the SNPs, a positive sign of *r* (or *r_v_*) strictly reflects both the coadaptation and compensatory epistatic models and a unilateral test can be performed with alternative hypothesis being “*r* (or *r_v_*) is significantly higher than 0”. Otherwise, the sign of *r* is not interpretable, but the p-value of the test can be computed on either side of the null distribution. Likewise, whether the ancestral/derived allele status of the SNPs is known or not, the sign of *cor_PC1_* (or *cor_PC1v_*) is not interpretable since *PC1* reflections imply identical ranking of individuals genotypes (or relatedness) in a given genomic region (see for instance (51)). R scripts to implement the statistical test based on *T*_*r*_, *T*_*rv*_, *T*_*corPC*1_ or *T*_*corPC*1*v*_, along with an example dataset, are available at https://github.com/leaboyrie/LD_corpc1.

### Simulation and LD-based detection power of epistatic selection in structured population

Simulations of the neutral evolution of two independent bi-allelic loci were carried out in order to evaluate the distribution of LD-based measures under the null hypothesis and their fit to a Student distribution τ_(*n*−2)_. In addition, two-locus epistatic selection was simulated in the same framework to evaluate the statistical power (1 – β) of these measures to detect two-locus epistatic selection given a type I error for the null hypothesis. Three selection models were simulated: the coadapted or compensatory epistatic selection model and the additive selection model (i.e. independent selection on the two loci; see table of fitness values in **S1 Table**). First, a sample from an ancestral population was simulated with the coalescence simulator SCRM (52). The sample consists of 1,000 haploid individuals with each four independent chromosomes of length L = 5 Mbp. The mutation (µ) and recombination (c) rates were fixed to classical values of 10^−9^ and 10^−8^ per base pair per generation, respectively, and the ancestral population size N_0_ fixed to 100,000, so that the ρ/θ ratio (i.e. 4N_0_cL/4N_0_µL) was equal to 10 and produced approximately 15,000 SNPs per chromosome in the sample, with a density of ∼ 1 SNP each 300 bp. The SCRM command used was *scrm nhap l –t theta –r rho l – p 8 > output.txt*. Second, the ancestral population of 1,000 haploid individuals was splitted into two isolated populations each of 250 diploid individuals, in order to generate population structure. Then, a forward-in-time simulation based on python language was carried using the SIMUPOP simulator (53, 54) under (i) different selection regimes: neutrality, coadapted or compensatory two-locus epistatic selection, or additive two-locus selection, and (ii) random or self-mating (with 95% selfing) scheme. Recombination and mutation probabilities were identical to SCRM. Simulation of each selection models were performed under complete recessivity, codominance or dominance of the mutations under selection. At generation 0, two SNPs were chosen approximately in the middle of two of the four different chromosomes, in order to follow allele frequencies and two-locus LD measures throughout 300 generations. At generation 100 in selection scenarios, the selection coefficients *s* was applied during 200 generations at these two SNPs in the two populations. Pilot studies suggested to use *s* = 0.10 to get adequate selection efficacy given the moderate simulated population size (N = 250 in each sub-population) and the computational cost to simulate small genomes with high SNP density. In addition, only coalescent simulations showing not too extreme derived allele frequencies at the two focal SNPs (i.e. between 0.25 and 0.75) were chosen to set up generation 0 of the forward-in-time simulation. Indeed, as suggested by previous works, standing variation on preselected mutations allows better efficacy of epistatic selection relative to genetic drift (10, 11, 13). Particularly under the coadaptation model, the fixation probability of the coadapted allelic combination highly depends on the initial allele frequencies, with low fixation probability when these mutations are in low frequencies at the onset of selection. Allele frequencies and the *r* and *r_v_* LD measures were calculated at the two SNP positions every 20 generations on the total population (500 diploid individuals distributed in two isolated populations). The *cor_PC1_* and *cor_PC1v_* measures were calculated on windows of 10 kbp (i.e. ± 5 kbp on either side of the focal SNP). For *r_v_* and *cor_PC1v_*, the kinship matrix was calculated at each time-point measure based on the simulated SNPs on the four chromosomes dataset, after filtering for a 5% Minor Allele Frequency (MAF). No MAF filtering was performed in the genomic regions of the two focal SNPs before calculating LD-based measures. We performed 1,000 simulations of each scenario described above, which allowed us to get the empirical distributions of all LD-based statistics.

### GWESS with SNP data in *Medicago truncatula* and human

GWESS was performed in *M. truncatula* using a raw dataset of 22,079,533 SNP markers identified on the eight chromosomes of the species by the Medicago HapMap Project on a collection of 262 accessions (see http://www.medicagohapmap.org/downloads/mt40). The collection has already been screened for GWAS of different traits (55–61) but also for GWSS (62–64). This highly self-mating species (95% selfing rate), originating from the Mediteranean basin, is structured in two major sub-populations, the Far West population (FW) concentrated on the West part under Atlantic influence, and the Circum population (C) that spreads over the rest of the Mediteranean basin (55, 56, 65, 66). Samples from the FW sub-population and from the C sub-population consist of 80 and 182 accessions, respectively. We used a bait approach for GWESS, in which *T*_*corPC*1_ and *T*_*corPC*1*v*_ were calculated for a given candidate gene, here *MtSUNN*, with gene identifier Medtr4g070970 in the genome version 4.0 – http://www.medicagogenome.org/ – (67) or MtrunA17Chr4g0035451 in the v5 – https://medicago.toulouse.inra.fr/MtrunA17r5.0-ANR/- (68), against each of the remaining 48,339 genes of the genome. For *PC1* calculation on each gene, imputed SNP data were required. Gene-based imputation was performed using the TASSEL software (69), where each missing base was imputed with the accession that shares the longest haplotype surrounding the base, on a window of 30 SNPs maximum (56). For calculation of *T*_*corPC*1*v*_, the kinship matrix *V* of the 262 individuals was estimated based on the genome-wide SNP dataset with MAF 5%.

In human, GWESS was performed by using the dataset of 494,764 SNPs (431,951 with a Minor Allele Frequency of 5%) with no missing data from the HGDP-CEPH Human Genome Diversity Panel on a world-wide sample (America, Asia, Europe, Middle East, North Africa, Subsaharian Africa and Oceania) of 940 individuals belonging to 57 populations from 23 countries (70, 71). The genome version (i.e. the gene positions) B36 was used for this analysis in order to fit with SNP positions in the HGDP-CEPH dataset, similarly to (72). We used a bait approach in which *T*_*r*_ and *T*_*rv*_ were calculated for SNPs located in or near *SLC24A5* and *EDAR* genes (chromosome 15 and 2, respectively), against each of the remaining 494,763 SNPs of the genome.

### Functional genetic validation of the relationship between *MtCLE02* and *MtSUNN in* Medicago truncatula

The Wild-Type *M. truncatula* A17 genotype from the Far West population was used in this study, as well as the *sunn-4* mutant allele derived from this genotype (73). Seeds were scarified by immersion in pure sulfuric acid for 3 min, rinsed six times with water, and sterilized for 20 min in Chlorofix (8.25 mg/L. Bayrol, France). After three washes with sterilized water, seeds were sown on 1% agar plates, and stratified for 3 days at 4°C in the dark. Germination was triggered by an overnight incubation at 24°C in the dark. To induce nodulation, a *Sinorhizobium medicae* WSM419 strain was used (74). Bacteria were grown overnight at 30°C on a Yeast Extract Broth (YEB) medium (75). For generating the *pAtUbi:MtCLE02* construct, the *MtCLE02* open reading frame was PCR amplified from *M. truncatula* genomic DNA using the Phusion High-Fidelity DNA polymerase (Thermo Scientific) and the 5’- <u>CACTCTGTGGTCTCAAATG</u>TAACAAACTTTTCTGATTCAT-3’ forward and 5’- <u>CACTTCGTGGTCTCAAAGC</u>TCAACCAATAAAAAACTATTAG-3’ reverse primers (underlines indicate Golden Gate adapter sites). Cloning was achieved using a Golden Gate strategy, as described in (76), with vectors generated by (77) and provided by the Engineering Nitrogen Symbiosis for Africa (ENSA) project (https://www.ensa.ac.uk/). The *pAtUbi:MtCLE02* single gene construct (promoter-gene-terminator) was cloned into a level 1 Golden Gate vector (EC47811, ampicillin resistant), which was then combined with a pNOS:Kanamycin cassette (EC15029), to allow *in planta* kanamycin selection of transformed roots, into a level 2 destination vector (EC50507). pAtUbi (EC15062) and 35S terminator (EC41414) cassettes were also provided by the ENSA project. A *pAtUbi:GUS* control vector was generated using the same strategy and the GUS cassette (EC75111). These binary vectors were introduced into the *Agrobacterium rhizogenes* ARqua1 strain to transform *M. truncatula* roots and generate composite plants as described in (78). Transformed roots were selected on kanamycin (50 µg/ml, Sigma), and plants with transgenic roots were then transferred in pots containing a mixture of perlite:sand (3:1) and the low nitrogen “i” medium (79), in a growth chamber (16h light at 150 µE intensity, 24°C, 60% relative air humidity). Plants were inoculated using a *S. medicae* bacterial suspension (OD_600nm_=0.05), and nodules were counted 14 days post inoculation (dpi). The *MtCLE02* transgene expression was validated by real time Reverse Transcriptase – Polymerase Chain Reaction (RT-PCR) as described in (80). In addition, *MtCLE02* expression was also analysed in a nodulation kinetic with the following timepoints: 0 (non-rhizobium inoculated roots), 1, 4, 8, 15 dpi (1 and 4 dpi corresponding to roots and 8 and 15 dpi to nodules). Primers used were as follows to amplify (i) *MtCLE02*: CLE02-Forward 5’-CAATGAATGTGAATGTTCTC-3’ and CLE02-Reverse 5’- TCAGGTCCATTAGGTACTCT-3’; (ii) *MtCLE13*: CLE13-Forward 5’- CCGAAGCCTTCTACAGAAACTACG-3’ and CLE13-Reverse 5’- TCTTGGTGGTGATCTTCCATTATGC-3’; and the *ACTIN11* reference gene: ACT11-Forward 5’-TGGCATCACTCAGTACCTTTCAACAG-3’ and ACT11-Reverse 5’- ACCCAAAGCATCAAATAATAAGTCAACC-3’.

## Acknowledgements

This work was supported by the “DeCoD” project funded by the French Agence Nationale de la Recherche (grant number ANR-16-CE20-0017-01). The PhD position of Léa Boyrie was funded by the “DeCoD” project. We thank the bioinformatics platform Toulouse Midi-Pyrenees (Genotoul). This work was performed in the LRSV (Toulouse, France), part of the “Laboratoire d’Excellence” (LABEX) entitled TULIP (grant number ANR-10-LABX-41). We thank Carole Laffont (IPS2, CNRS, Gif-sur-Yvette, France) for providing results about *MtCLE13* expression. Work in the Florian Frugier laboratory has benefited from a French State grant (Saclay Plant Sciences, grant number ANR-17-EUR-0007, EUR SPS-GSR) and ANR grant PSYCHE (grant number ANR-16-CE20-0009-01). We thank Pierre-Marc Delaux for useful criticisms and comments on the manuscript.

---

**Table S1: Fitness for genotypes at two loci (A and B) in epistatic selection models (COAD and COMP) and an additive (ADD) selection model, for recessive, dominant and codominant mutations (derived alleles *a* and *b*).**

**Figure S1. Evolution of fixation index *F_ST_* and inbreeding coefficient *F_IS_* in the simulated data over 300 generations and example of kinship matrices at the outcome of a simulation.**

Evolution of *F_ST_* and *F_IS_* coefficients in simulations of self-mating (**A**) and random-mating (**B**) systems, calculated from neutral models. Heatmaps of kinship matrix (generation 300) showing the distribution of genomic relatedness among individuals in presence of population structure and contrasted reproductive systems: self-mating (**C**) and random-mating (**D**).

**Figure S2. Co-fixation rate of mutant (*ab*) and derived (*AB*) haplotypes at simulated loci.**

Co-fixation rate of mutant (*ab)* and ancestral (*AB*) haplotypes over generations showing selection efficiency in self-mating (**A** – **C** – **E**) and in random mating model (**B** – **D** -**F**). Co-fixation dynamic is depicted for recessive (**A**-**B**), codominant (**C** – **D**) and dominant (**E** – **F**) mutations. Note that alleles *A*/*a* and *B*/*b* are the alleles of SNPs A and B under epistatic selection, which are located in the middle of two of the four simulated chromosomes.

**Figure S3. Evolution of inter-locus epistatic selection detected with linkage disequilibrium on simulated data.**

Mutations under epistatic selection are recessive (**A, B, C, D**) or dominant (**E, F, G, H**). Inter-locus LD in self-mating simulation model is calculated on SNP-based scale with *r* (*r_v_*) (**A, E**) and on window-based scale with *cor_PC1_* (*cor_PC1v_*) (**B, F**). Inter-locus LD in random mating simulation model is calculated on SNP-based scale (**C, G**) and on window-based scale (**D, H**).

**Figure S4. Expression of *MtCLE02* in a nodulation kinetic and in overexpressing roots.**

**(A)** Real time RT-PCR expression analysis of *MtCLE02* in a nodulation kinetic, 0 (non-inoculated roots), 1, 4, 8 or 15 days post rhizobium inoculation (dpi). 1 and 4 dpi correspond to roots, and 8 and 15 dpi to nodules. **(B)** Real time RT-PCR expression analysis of *MtCLE13*, a CLE signaling peptide encoding gene previously linked to symbiotic nodulation, in the same conditions as described in **A**. **(C)** Real time RT-PCR expression analysis of *MtCLE02* in roots overexpressing *MtCLE02* (Ubi:CLE02) or the *GUS* control (Ubi:GUS), which symbiotic nodulation phenotype is analyzed in Figure 4. Two independent transgenic roots were analyzed for each genotype.

**Figure S5. LD distribution between the bait SNPs of *SLC24A5* and *EDAR* genes and all other HGDP-CEPH SNPs in the <u>Central South Asia</u> human population samples (n=192).**

LD between SNP_15_46172199 (*SLC24A5)* and SNP_2_108973688 (*EDAR*), respectively, and all other SNPs of the genome is tested using *T*_*r*_ (**A, B**) or *T*_*rv*_ (**C, D**), which includes the kinship matrix. The *x*-axis corresponds to SNP positions spanning the 22 human autosomes, each point corresponds to a SNP and the black points depict SNPs at candidate genes in epistatic selection with one SNP at the bait gene (vertical dotted line) in each figure. The y-axis is the −log_10_(p-value) of the test of the correlation coefficient. Plots at the top left of each figure show distribution of LD between each bait SNP and all other SNPs of the genome. LD value between bait SNPs of *SLC24A5* and the target SNP of *EDAR* (respectively bait SNPs of *EDAR* and target SNP of *SLC24A5*) is represented by an arrow.

**Figure S6. LD distribution between the bait SNPs of *SLC24A5* and *EDAR* genes and all other HGDP-CEPH SNPs in the <u>East Asian</u> human population samples (n=242).**

**Figure S7. LD distribution between the bait SNPs of *SLC24A5* and *EDAR* genes and all other HGDP-CEPH SNPs in the <u>Subsaharian Africa</u> human population samples (n=105).**

**Figure S8. LD distribution between the bait SNPs of *SLC24A5* and *EDAR* genes and all other HGDP-CEPH SNPs in the <u>Middle East</u> human population samples (n=134).**

**Figure S9. LD distribution between the bait SNPs of *SLC24A5* and *EDAR* genes and all other HGDP-CEPH SNPs in the <u>European</u> human population samples (n=158).**

**Figure S10. LD distribution between the bait SNPs of *SLC24A5* and *EDAR* genes and all other HGDP-CEPH SNPs in the <u>America</u> human population samples (n=64).**

For Figures S6 to S10, the legend is the same as in Figure S5.

**Figure S11. Distribution of LD between SNPs at *SLC24A5* and *EDAR* genes.**

Average and standard error of LD significances based on *T*_*r*_ statistics between SNPs of *SLC24A5* (SNP 15_46172199, SNP 15_46179457) and *EDAR* (SNP 2_108962124, SNP 2_108973688, SNP 2_108982808) within each human sub-population.

