## Supplementary Information Files for "A fast linkage disequilibrium-based statistical test for Genome-Wide Epistatic Selection Scans in structured populations"

Boyrie et al.

Submission to *PLOS Genetics*

**Table S1. Fitness for genotypes at two loci (A and B) in epistatic selection models (COAD and COMP) and an additive (ADD) selection model, for recessive, dominant and codominant mutations (derived alleles *a* and *b*).**

| | | COAD | | | COMP | | | ADD | | |
|---|---|---|---|---|---|---|---|---|---|---|
| | | $BB$ | $Bb$ | $bb$ | $BB$ | $Bb$ | $bb$ | $BB$ | $Bb$ | $bb$ |
| *recessive* | $AA$ | 1 | 1 | 1 | 1 | 1 | 1-2s | 1 | 1 | 1+s |
| | $Aa$ | 1 | 1 | 1 | 1 | 1 | 1-2s | 1 | 1 | 1+s |
| | $aa$ | 1 | 1 | 1+2s | 1-2s | 1-2s | 1 | 1+s | 1+s | 1+2s |
| *codominant* | $AA$ | 1 | 1 | 1 | 1 | 1-s | 1-2s | 1 | 1+s/2 | 1+s |
| | $Aa$ | 1 | 1+s | 1+s | 1-s | 1-s | 1-s | 1+s/2 | 1+s | 1+1.5s |
| | $aa$ | 1 | 1+s | 1+2s | 1-2s | 1-s | 1 | 1+s | 1+1.5s | 1+2s |
| *dominant* | $AA$ | 1 | 1 | 1 | 1 | 1-2s | 1-2s | 1 | 1+s | 1+s |
| | $Aa$ | 1 | 1+2s | 1+2s | 1-2s | 1 | 1 | 1+s | 1+2s | 1+2s |
| | $aa$ | 1 | 1+2s | 1+2s | 1-2s | 1 | 1 | 1+s | 1+2s | 1+2s |

**Figure S1. Evolution of fixation index $F_{ST}$ and inbreeding coefficient $F_{IS}$ in the simulated data over 300 generations and example of kinship matrices at the outcome of a simulation.**

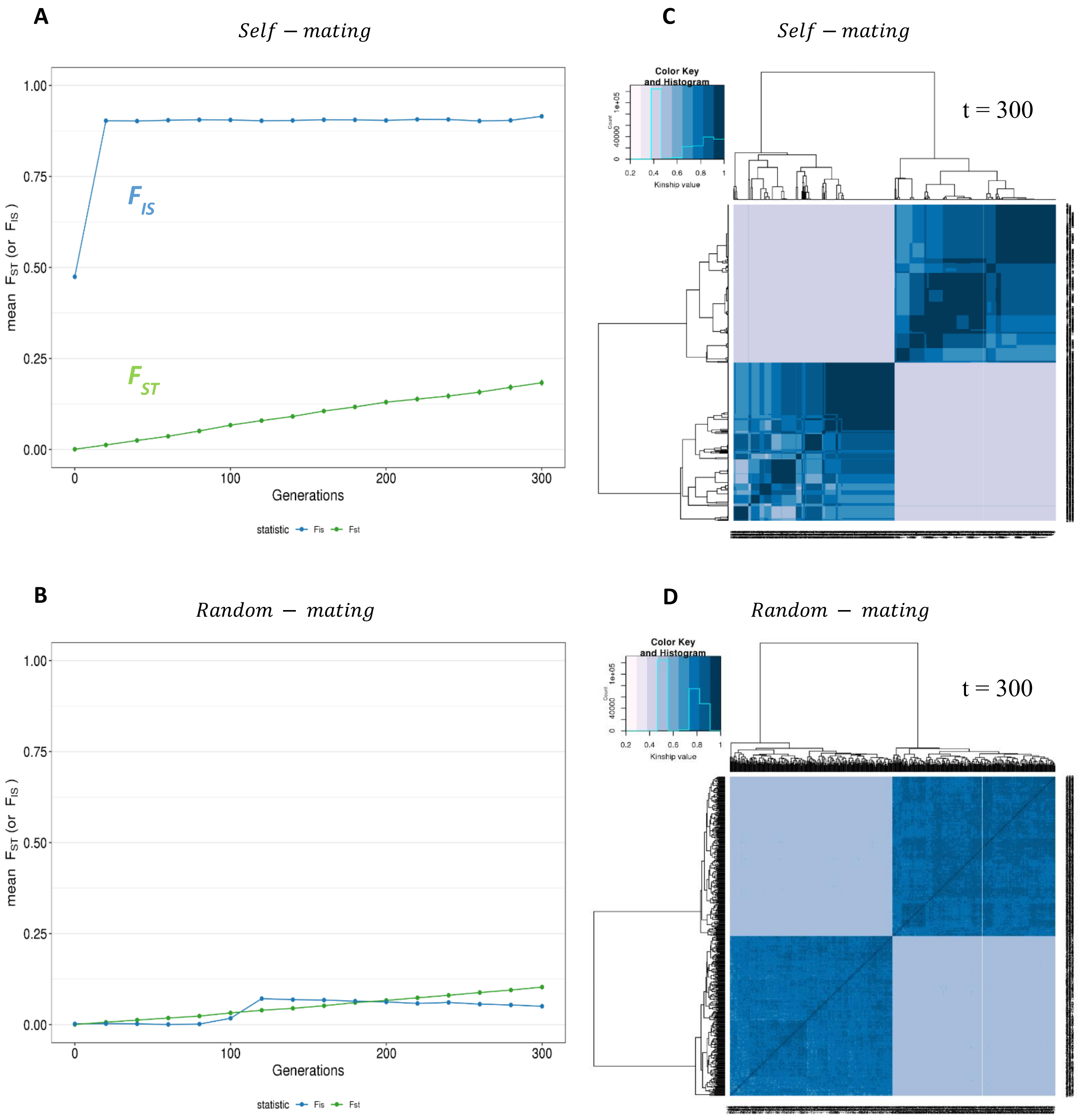

**Figure S2. Co-fixation rate of mutant (*ab*) and derived (*AB*) haplotypes at simulated loci.**

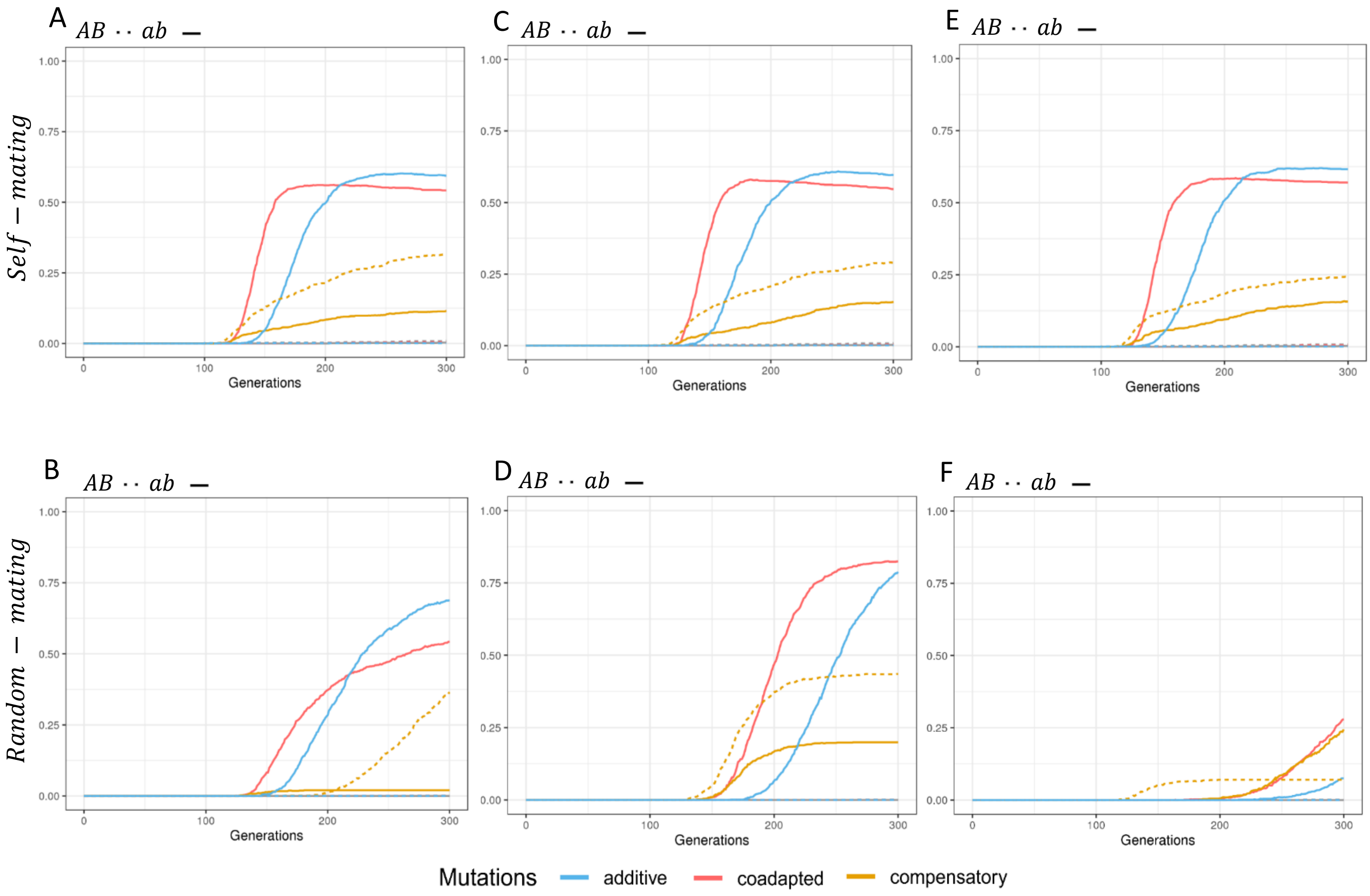

**Figure S3. Evolution of inter-locus epistatic selection detected with linkage disequilibrium on simulated data.**

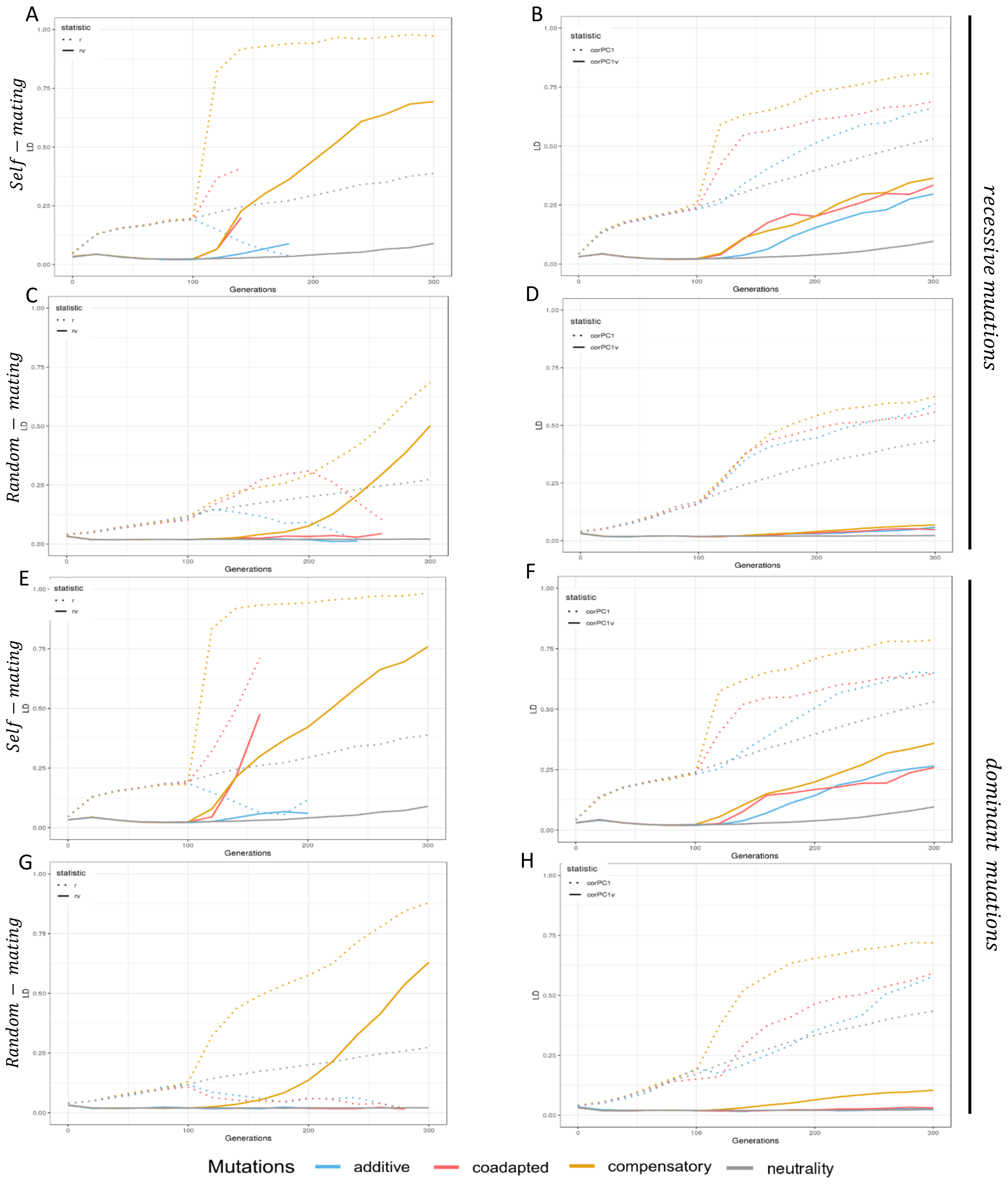

**Figure S4. Expression of *MtCLE02* in a nodulation kinetic and in overexpressing roots.**

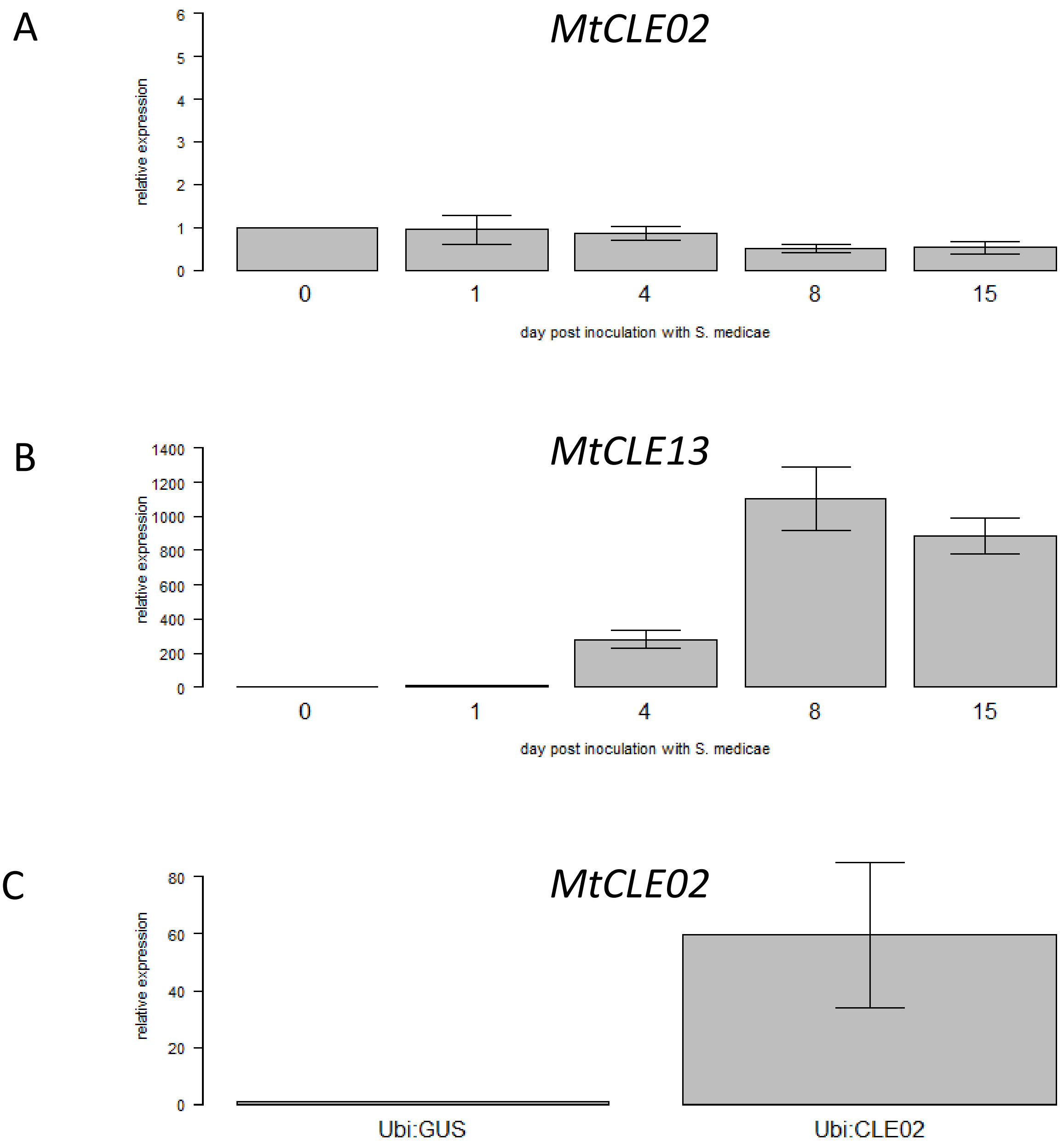

**Figure S5. LD distribution between the bait SNPs of *SLC24A5* and *EDAR* genes and all other HGDP-CEPH SNPs in the Central South Asia human population samples (n=192).**

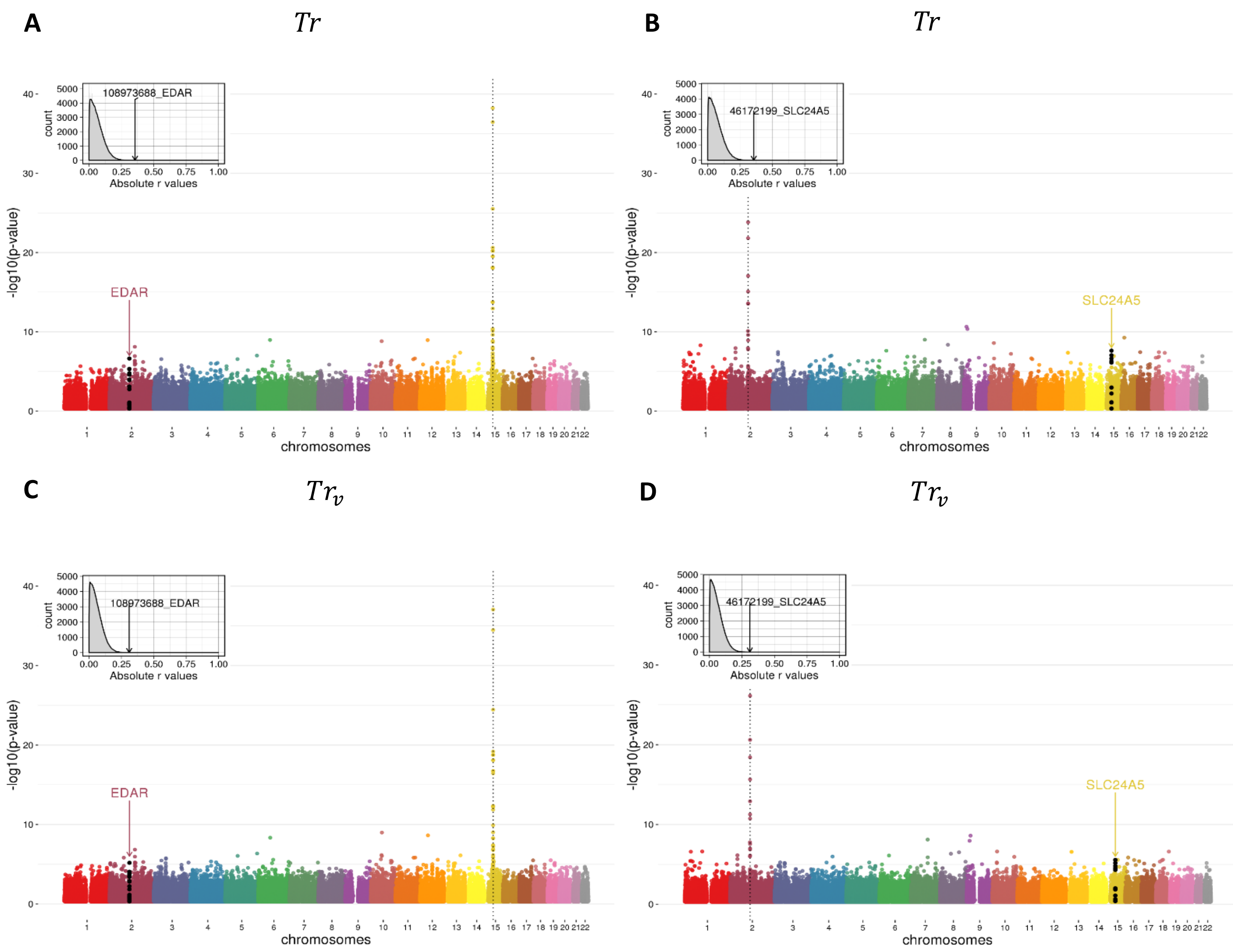

**Figure S6. LD distribution between the bait SNPs of *SLC24A5* and *EDAR* genes and all other HGDP-CEPH SNPs in the East Asian human population samples (n=242).**

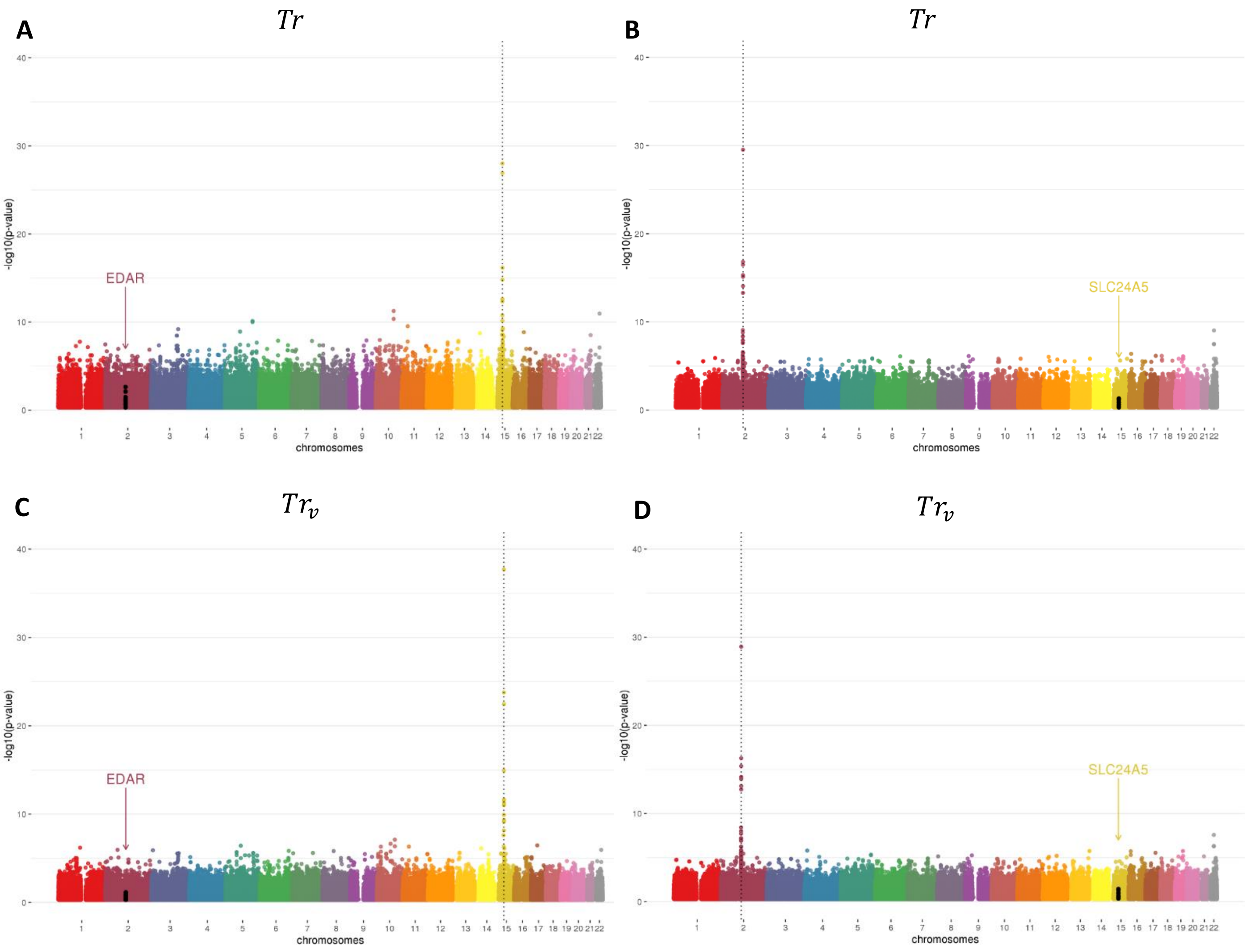

**Figure S7. LD distribution between the bait SNPs of *SLC24A5* and *EDAR* genes and all other HGDP-CEPH SNPs in the <u>Subsaharian Africa</u> human population samples (n=105).**

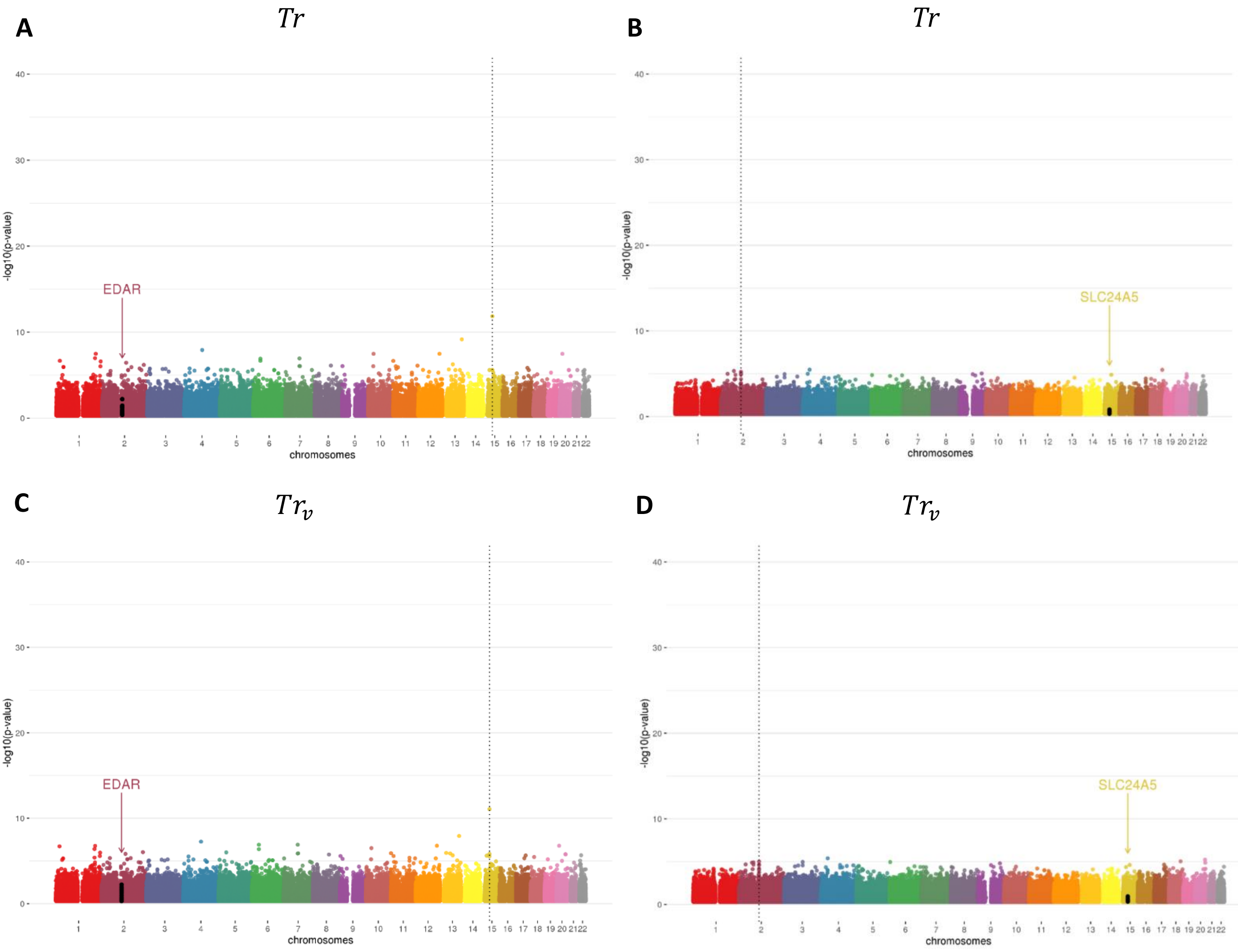

**Figure S8. LD distribution between the bait SNPs of *SLC24A5* and *EDAR* genes and all other HGDP-CEPH SNPs in the <u>Middle East</u> human population samples (n=134).**

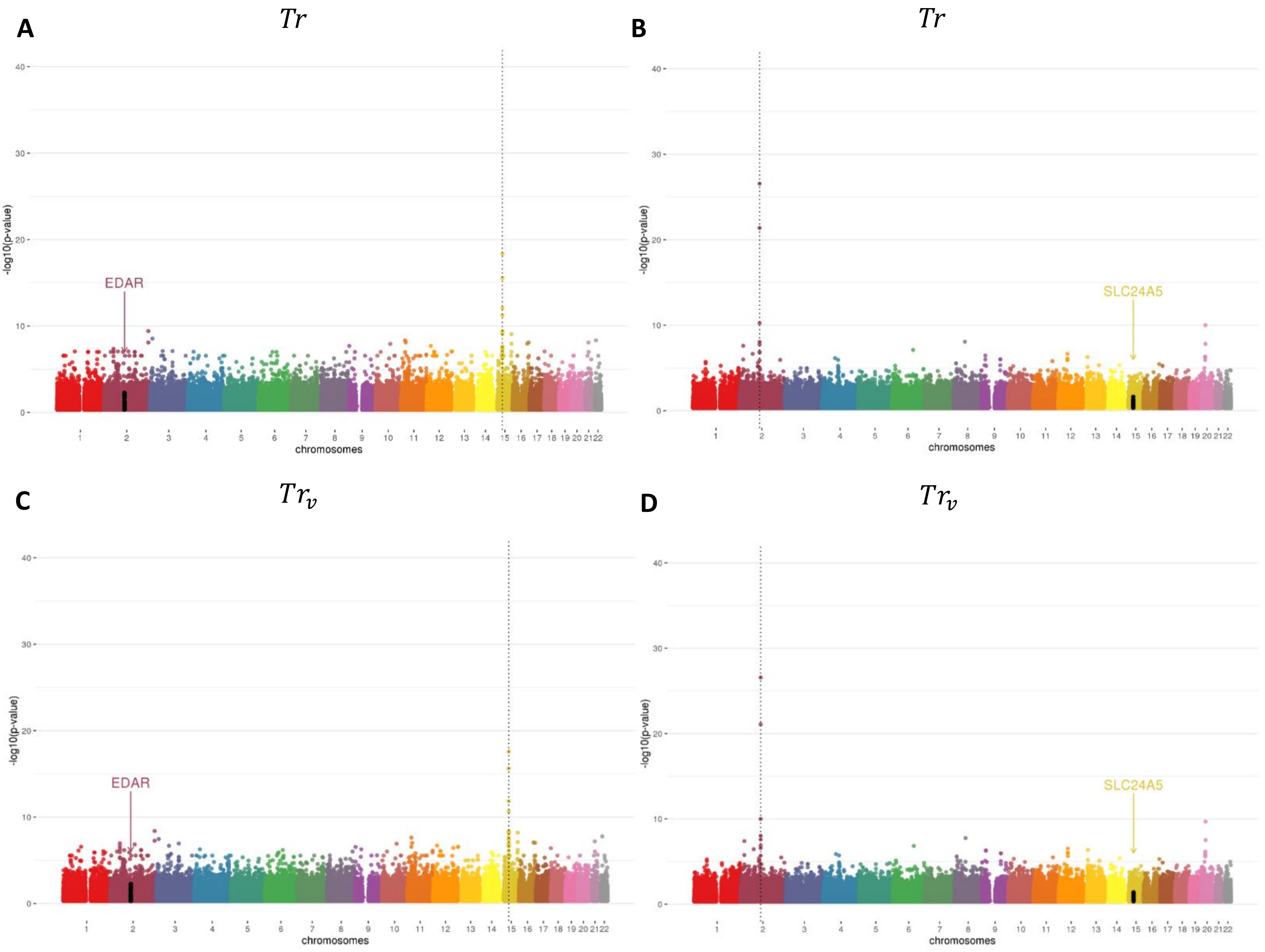

**Figure S9. LD distribution between the bait SNPs of *SLC24A5* and *EDAR* genes and all other HGDP-CEPH SNPs in the <u>European</u> human population samples (n=158).**

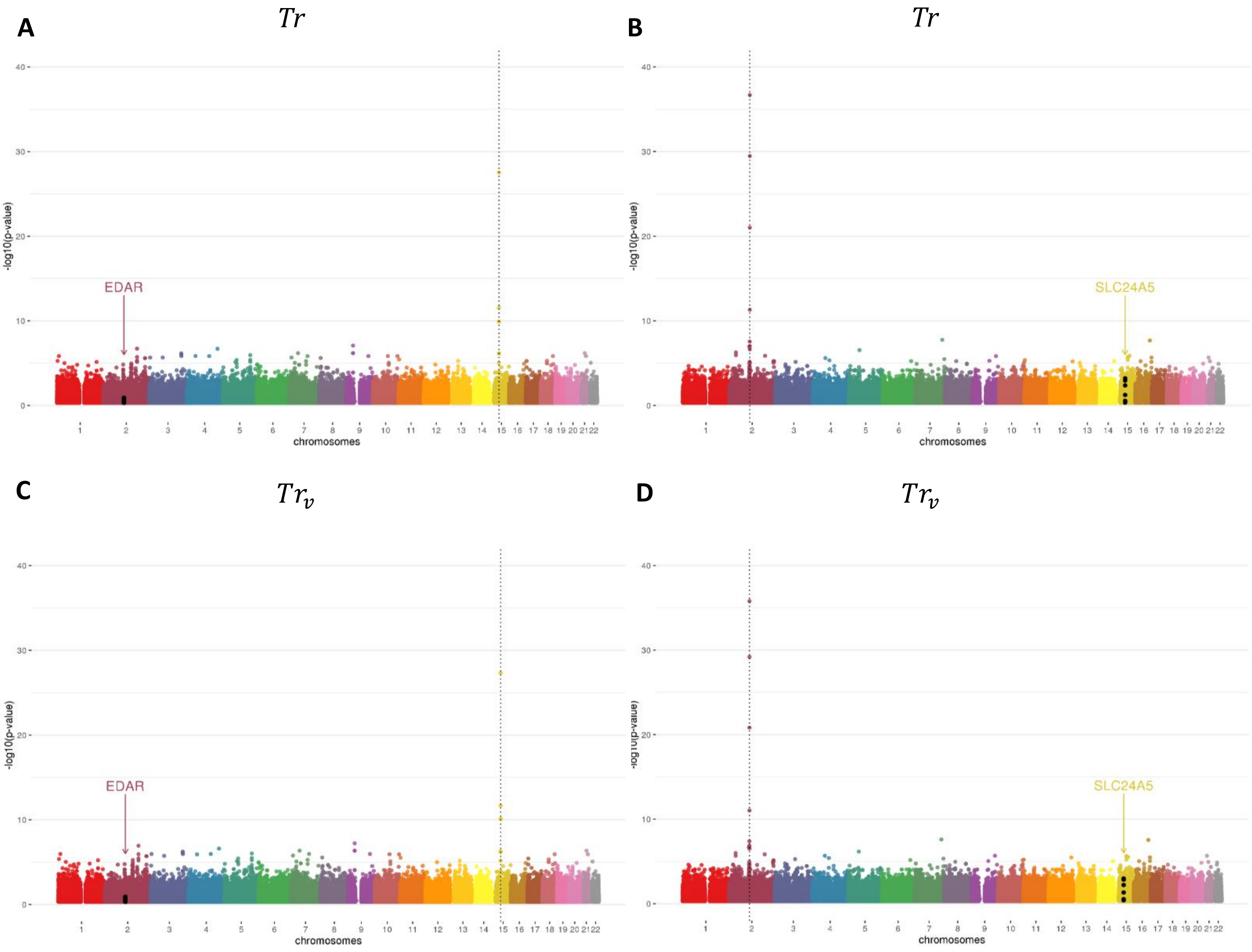

**Figure S10. LD distribution between the bait SNPs of *SLC24A5* and *EDAR* genes and all other HGDP-CEPH SNPs in the <u>America</u> human population samples (n=64).**

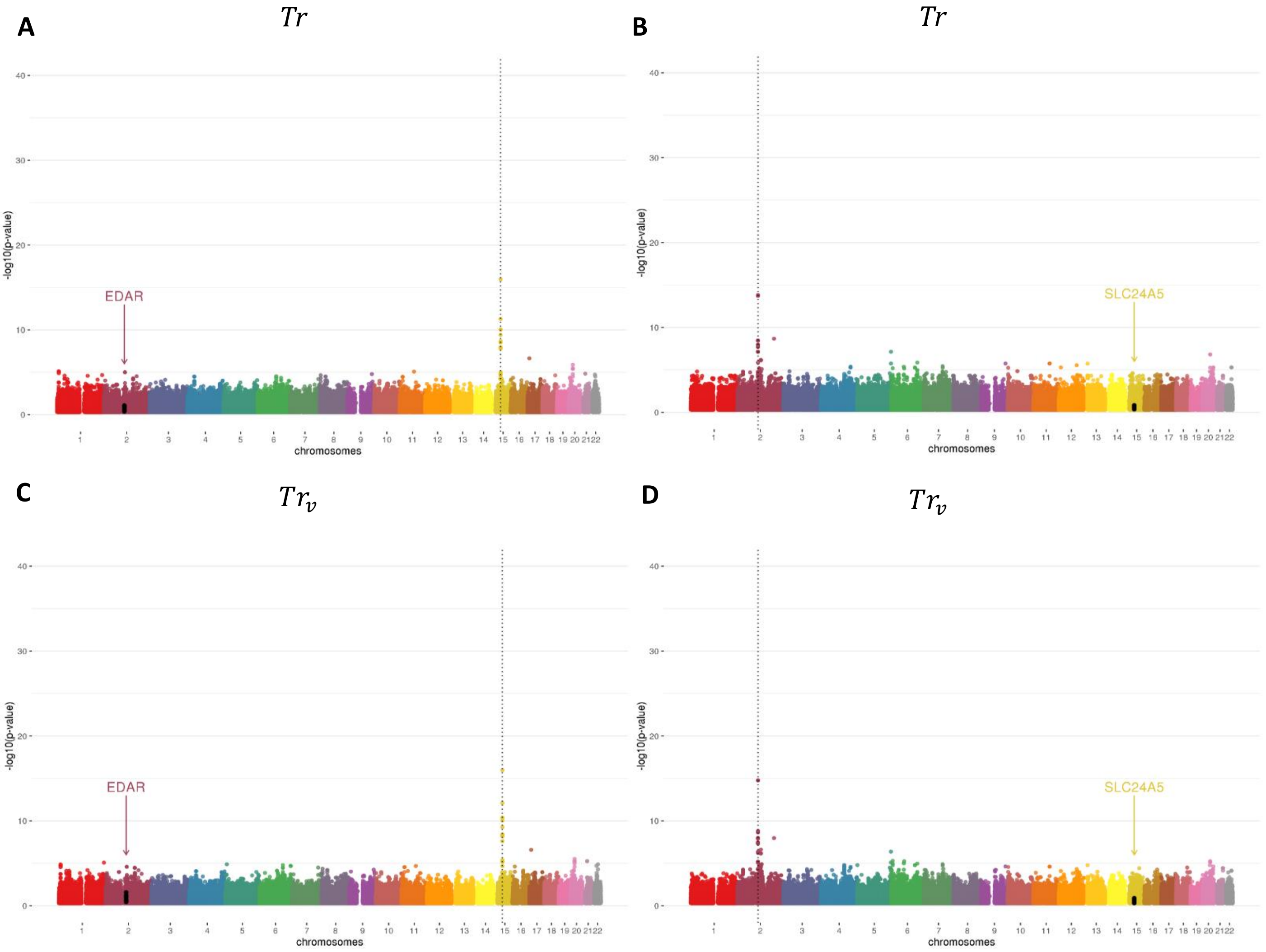

**Figure S11. Distribution of LD between SNPs at *SLC24A5* and *EDAR* genes.**

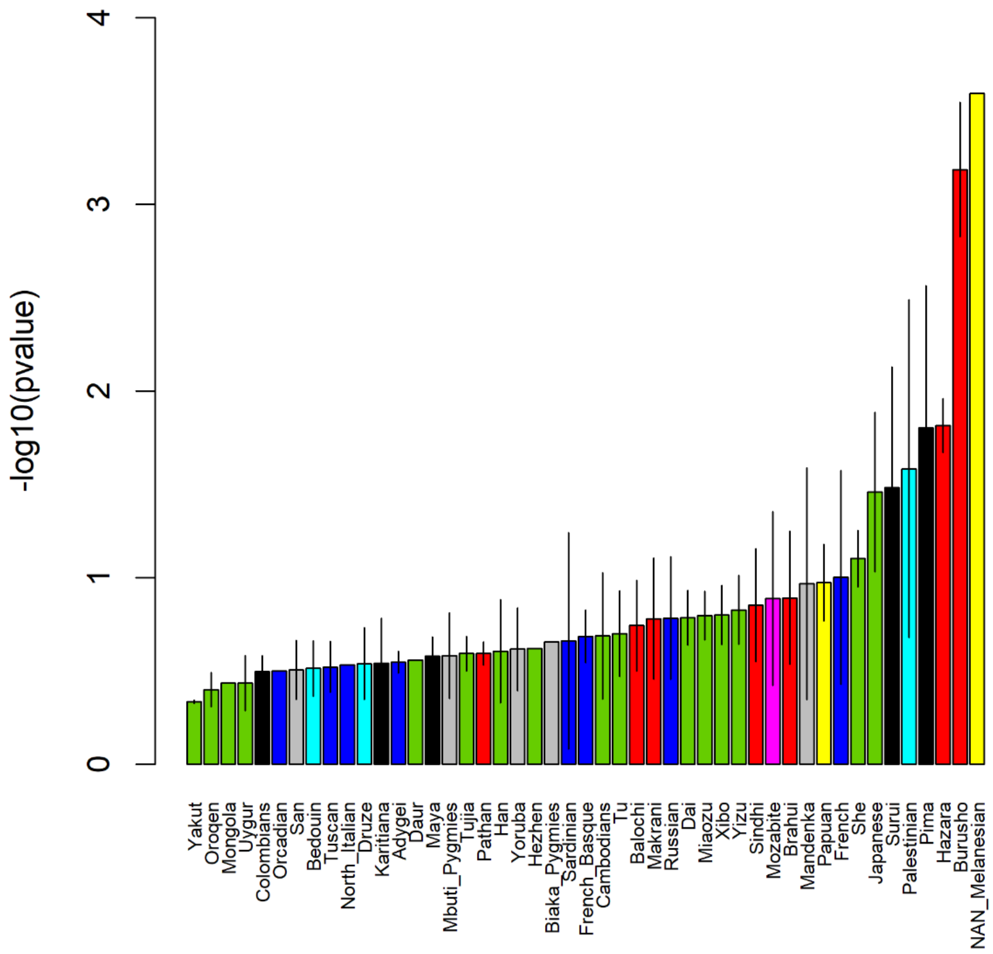
